# Visualizing homology search in living cells

**DOI:** 10.1101/2025.03.01.640932

**Authors:** Anoek Friskes, Melanie Snoek, Roel Oldenkamp, Bram van den Broek, Leila Nahidiazar, Lisa Koob, Amalie E. Dick, Marjolijn Mertz, Rolf Harkes, Benjamin D. Rowland, Kees Jalink, René H. Medema

## Abstract

Homologous recombination (HR) is a DNA repair process that requires a homologous sequence for the repair of DNA double-stranded breaks. The search for homology is the rate limiting step for HR, but the dynamics of homology search *in vivo* are currently poorly understood. This is largely due to a lack of tools to visualize homology search in action in living human cells. Here, we show that RAD51 and MND1, two proteins operating in homology search, become visible in long filamentous structures several hours after double-stranded break formation. Using GFP-MND1 we can visualize these filaments in living human cells and find that they are highly dynamic, explore the nuclear space, and can resolve over time. We show that the resolution of these filaments depends on RAD54L, known for its role in RAD51-driven homology search and strand-invasion. In addition, we find that loss of cohesin also inhibits the resolution of these filaments, in accordance with a role for cohesin in HR. Thus, our data suggest that these filaments are visible intermediates of an active DNA repair process, and that MND1-GFP can be used as a tool to study the dynamics of homology search in living human cells.

## Introduction

Error-free repair of broken DNA is essential for genome maintenance. One of the principal pathways employed by cells to faithfully repair DNA double-strand breaks (DSBs) is homologous recombination (HR) ^1^. This process relies on the use of an intact homologous DNA sequence as a template for accurate repair ^2–4^. In human somatic cells, HR is (most) active in S/G2 cells, when a sister chromatid is available and usually provides the repair template. If the sister chromatid cannot be used, sequences on the homologous chromosome can be used as templates instead, resulting in interhomolog recombination ^5^. In such cases, the search for the correct repair template in the vast and crowded nuclear space can become a challenge. This challenge comes with the risk of selecting an incorrect donor sequence, thereby promoting genomic instability ^5,6^.

Homology sampling is mediated by a nucleoprotein filament, consisting of the recombinase protein RecA (bacteria) or RAD51 (eukaryotes) bound to ssDNA ^7^. *In vitro* studies have shown that RecA/RAD51 can form nucleoprotein filaments on ssDNA ^8^, and guide the coated ssDNA along the double-stranded DNA to identify the homologous sequence ^9^. To engage in HR, the 5’ends on either side of the DNA double-stranded break are first resected by nucleases such as MRE11, EXO1, and DNA2. Once ssDNA is generated, it is rapidly coated and stabilized by the ssDNA-binding protein complex RPA (replication protein A) which is subsequently replaced by the recombinase RAD51 ^10^. RAD51 nucleoprotein filament formation is an important step in HR, and a failure to replace RPA with RAD51 can create intermediates that cannot be properly processed and that can become detrimental to cells ^11^.

Already decades ago, studies conducted in bacteria showed that RecA forms elongated structures associated with homology search and strand invasion at sites of DSBs ^12–15^. In line with this, various subsequent genetic, *in vitro*, structural, and molecular studies indicated that RAD51 can also form nucleoprotein filaments ^9,16–21^. This concept was more recently supported by live-cell imaging of endogenously GFP-tagged Rad51 in yeast, demonstrating highly dynamic nucleoprotein filaments undergoing rounds of extension and compaction ^22^. These filaments can span the entire yeast nucleus and establish contact with the template sequence. Thus far, real-time dynamics of homology search in human cells have not been visualized.

In the current study, we use quantitative 3D super-resolution imaging of fixed cells, complemented with high-resolution live-cell imaging, and show that puncta of RAD51, as well as MND1, that form early after double-stranded break formation, stretch into filamentous structures later during the repair process. We show that these filaments dynamically explore the nuclear space and resolve over time. We show that their resolution depends on RAD54L and cohesin, both proteins with a previously established role in HR. We propose that the formation of these structures reflects a crucial step in homologous recombination and that GFP-MND1 represents a useful tool to study the dynamics of homology search in human cells.

## Results

### RAD51 forms extended filaments at sites of DNA damage

It was previously shown that RAD51 accumulates at DNA lesions following irradiation-induced DNA damage. While most studies report the formation of (spherical) RAD51 foci, some have shown that RAD51 filaments can also be observed at higher resolution ^23,24^. Elegant single-molecule localization microscopy experiments in HeLa cells have demonstrated that RAD51 clusters at DNA damage foci and progressively extends into filaments over time ^23^. These extended filaments undergo topographical changes and reach a maximal length ∼3-5 hr after damage induction, suggesting that they represent RAD51 nucleoprotein filaments engaged in homology search ^23^. To explore RAD51 nucleoprotein filament dynamics in non-transformed human cells, we used immortalized RPE-1 cells. In these cells, RAD51 foci formation is observed within 2 hr after damage, and remains high at 4 and 6 hr after damage induction, while at 16 hr most of the RAD51 and γH2AX foci are resolved (Fig.1A-C). γH2AX marks sites of DNA damage, and therefore these data imply that a significant part of the breaks that engage in RAD51-dependent repair are repaired between 6 and 16 hr. At 16 hr after IR, a fraction of RAD51 staining appeared as small dots, while others appeared as filaments of variable lengths and topography (Fig.1D). For simplicity, we refer to the dot-like foci as “RAD51 puncta”, and the extended assemblies as “RAD51 filaments” throughout this manuscript. RAD51 filaments frequently extend away from the γH2AX-positive core of the DNA damage focus (Fig.1E, Suppl. Fig.1A). This implies that the RAD51 filaments we observe are associated with a broken DNA end (Fig.1E, Suppl. Fig.1A). In addition, the lack of γH2AX-staining on the filaments suggests that this part of the nucleoprotein filament is free of nucleosomes, or that γH2AX has been removed. The dual appearance of RAD51 in puncta and filaments is consistent across multiple cell lines and seems independent of p53 status (i.e., wild-type or mutant p53), although the length of the filaments appears to vary (Suppl. Fig.1B). RAD51 filaments formed in S-phase as well as in G2 phase (Suppl. Fig.1C). When comparing filament topography at 6 versus 16 hr after irradiation, we noted that filaments were longer and less spherical at 16 hr (Fig.1F), suggesting that RAD51 filaments that are not resolved continue to grow in size.

**Fig 1:**
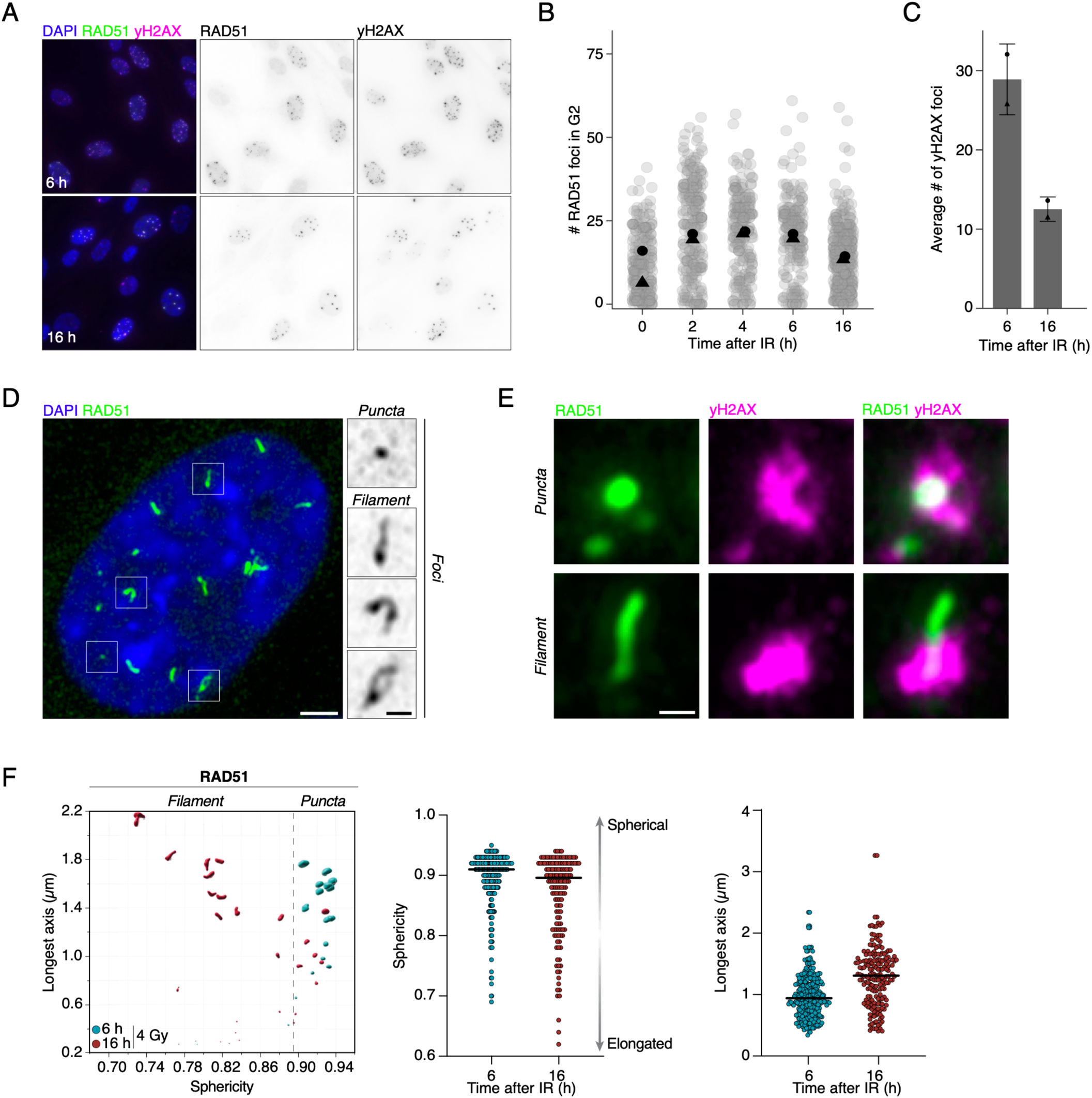
RAD51 forms extended filaments at sites of DNA damage. A) Wide-field microscopy examples of RPE-1 HALO-53BP2=1 cells exposed to irradiation (4 Gy, 6 hours and 16 hours), immunostained with γH2AX (magenta) and RAD51 (green). B-C) Quantification of the amount of RAD51 (B) or γH2AX (C) followed indicated hours after irradiation (4 Gy) as shown in (A). N=2, indicated by the dots. D) Immunofluorescence staining of RAD51 in RPE-1 wild-type cells resolved by super-resolution as filament structure throughout the nuclear volume (blue; DAPI). Cells were irradiated with 4 Gy and fixed 16 hours later. Scale bar, 2 μm. Insets, 0.5 μm. E) Immunofluorescence staining of RAD51 (green) puncta and filaments and yH2AX (magenta) in RPE-1 wild-type cells resolved by super-resolution. Cells were irradiated with 4 Gy and fixed 16 hours later. Scale bar, 1 μm. F) Example plot of the sphericity against the longest axis of RAD51 foci measured in 3D, 6 or 16 hours after irradiation (4 Gy) of one cell. The plot of the sphericity and longest axis of all RAD51 foci measured in many cells. The higher the value for sphericity, the more spherical (puncta) a focus is, the lower the value, the more elongated the focus is. Data were obtained through Imaris.

RAD51 filaments can adopt a variety of confirmations (Fig.2A) that closely resemble the RAD51 filaments observed by single-molecule localization microscopy in HeLa cells and yeast ^22,23^. We could discern distinct filament structures which we classified as small rods, rods, and coiled/branched filaments (Fig.2A), and we employed Ilastik Machine learning to quantify these subclasses (Fig. 2B). When comparing the combined frequency of these three subclasses at 6 and 16 hr, we observed a clear shift in the puncta/filament ratio, so that more than half of all the remaining RAD51 foci at 16 hr appeared as filaments (Fig.2C). There here is a clear increase in filament length (Suppl. Fig.2A), filament area (Suppl. Fig.2A) and complexity (Fig.2D) at 16 hr after damage. The average longest path of RAD51 filaments for instance increased from ∼500nm at 6 hours to ∼900nm at 16 hours post-damage (Suppl. Fig.2A). As these lengths align well with reported distances between sister chromatids in G2 ^25,26^, we speculate that the observed RAD51 filament represents the RAD51-coated nucleoprotein filament that engages in homology search to identify the proper template on the sister chromatid or the homologous chromosome. We note that at 16 hr, a small part of the initial DNA damage foci remain unresolved (Fig.1B,C). The remaining RAD51 filaments presumably failed to identify a suitable repair template and continue to grow because of ongoing resection, explaining the increase in length and complexity seen at 16 hr. Taken together, these RAD51 filaments we observe in human cells could be relevant representations of RAD51-nucleoprotein filaments that engage in homology search, like the proposed role for Rad51 filaments in yeast.

**Fig 2:**
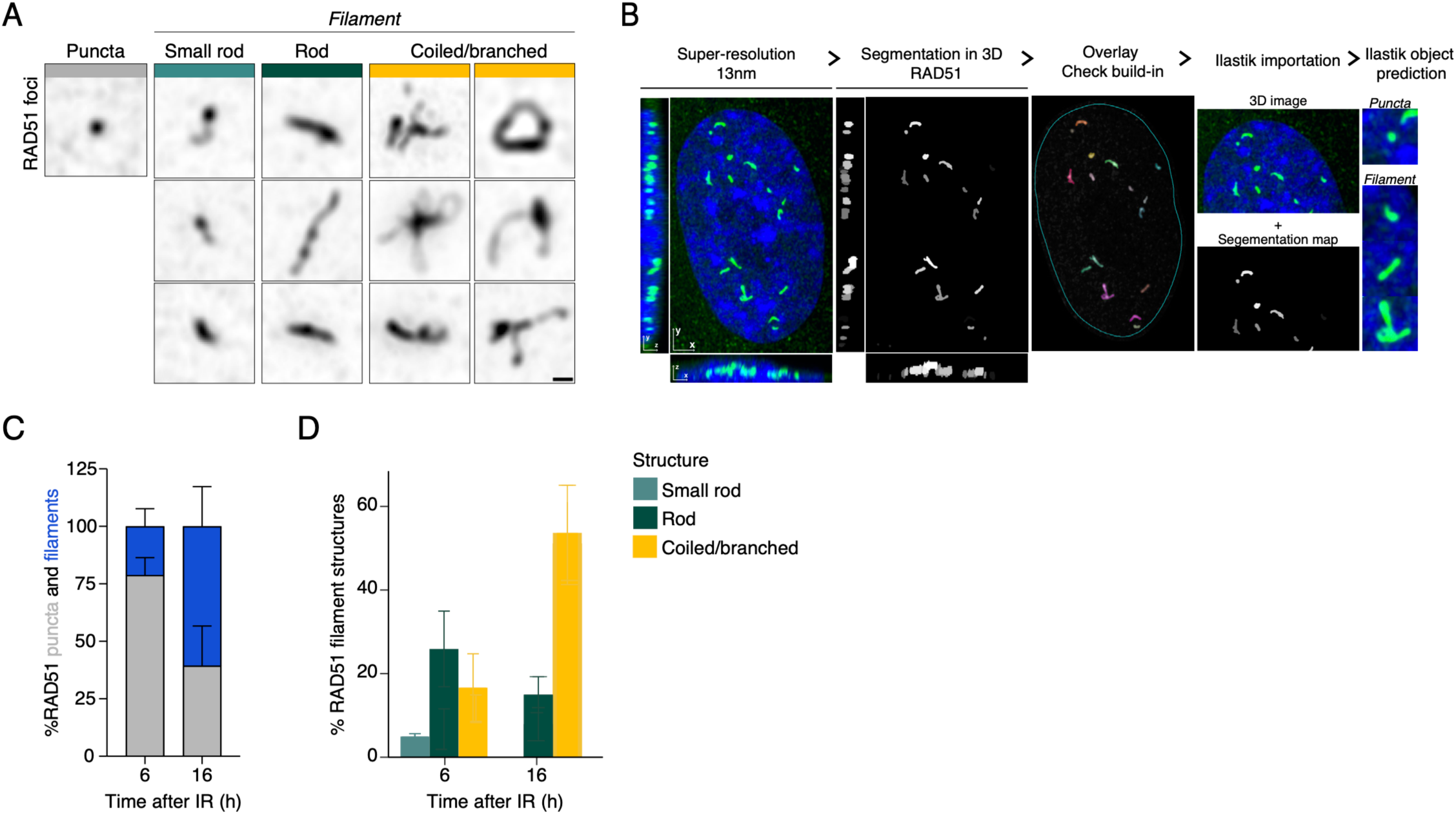
Super-resolution imaging of RAD51 reveals their intracellular localization. A) Super-resolution microscopy examples of various RAD51 foci assemblies. Puncta, small rods, rods and coiled/branches categories were made of different shapes (16 hours, 4 Gy): scale bar, 0.5 μm. B) Columns 1 and 2: Machine learning–assisted segmentation of the nucleus and RAD51 structures. Lateral (top) and orthogonal (bottom) cross sections of 3D image stacks are shown. Column 3: Maximum intensity projection of both the original super-resolution image and the segmentation which closely resembles the structures. Column 4: Both the original 3D super-resolution image and segmentation map (in 3D) were imported into Ilastik. Column 5: Ilastik was trained to predict either puncta or filaments. C) Average percentages of RAD51 puncta and filaments at 6, 16 hours after damage. Data were obtained through Ilastik. N=2. D) Ilastik analysis of various RAD51 foci assemblies of RPE1 cells. Puncta, small rods, rods and coiled/branches categories were made of different shapes as shown in (A) (6/16 hours, 4 Gy): N=2.

### RAD54L is required for the resolution of RAD51 filaments

Several co-factors have been shown to control the formation, stabilization and strand invasion of RAD51-nucleoprotein filaments. RAD54L is a member of the SWI2/SNF2 family of DNA-dependent ATPases that acts at several stages during homology search. It was shown to stabilize the RAD51 nucleoprotein filament, to be required for D-loop formation, and to remove RAD51 from heteroduplex DNA after strand invasion ^27–30^. Indeed, depletion of RAD54L from RPE-1 cells resulted in a delay of RAD51-dependent repair, as evidenced by an increase in residual RAD51 foci at the 6 and 16 hr time points (Fig 3A). Consistently, HAP1 cells in which RAD54L was knocked out (HAP1ΔRAD54L) also displayed a mild increase in sensitivity to irradiation (Suppl. Fig.3A,B). To test if RAD54L affects RAD51 filament dynamics, we generated RAD54L knock-out clones in RPE-1 background (RPE-1ΔRAD54L) and imaged RAD51 foci (Suppl. Fig. 3C). Notably, ΔRAD54L cells already displayed RAD51 filaments in the absence of irradiation (Fig.3B). At 6 hr after irradiation the percentage of RAD51 filaments was increased in RAD54L knock-out cells (Fig.3C). Also, absence of RAD54L resulted in longer filaments that covered more area (Fig.3D-F). Similar effects were observed in HAP1ΔRAD54L cells (Suppl. Fig.3D). Thus, when RAD51 filament formation is allowed, but RAD51-dependent repair is perturbed by the depletion of RAD54L, more filaments accumulate and become longer and more complex (Fig.3G). These data further support the notion that RAD51 filaments that extend away from the core of a DNA damage focus and grow in length over time might be a visual representation of the homology search process.

**Fig 3:**
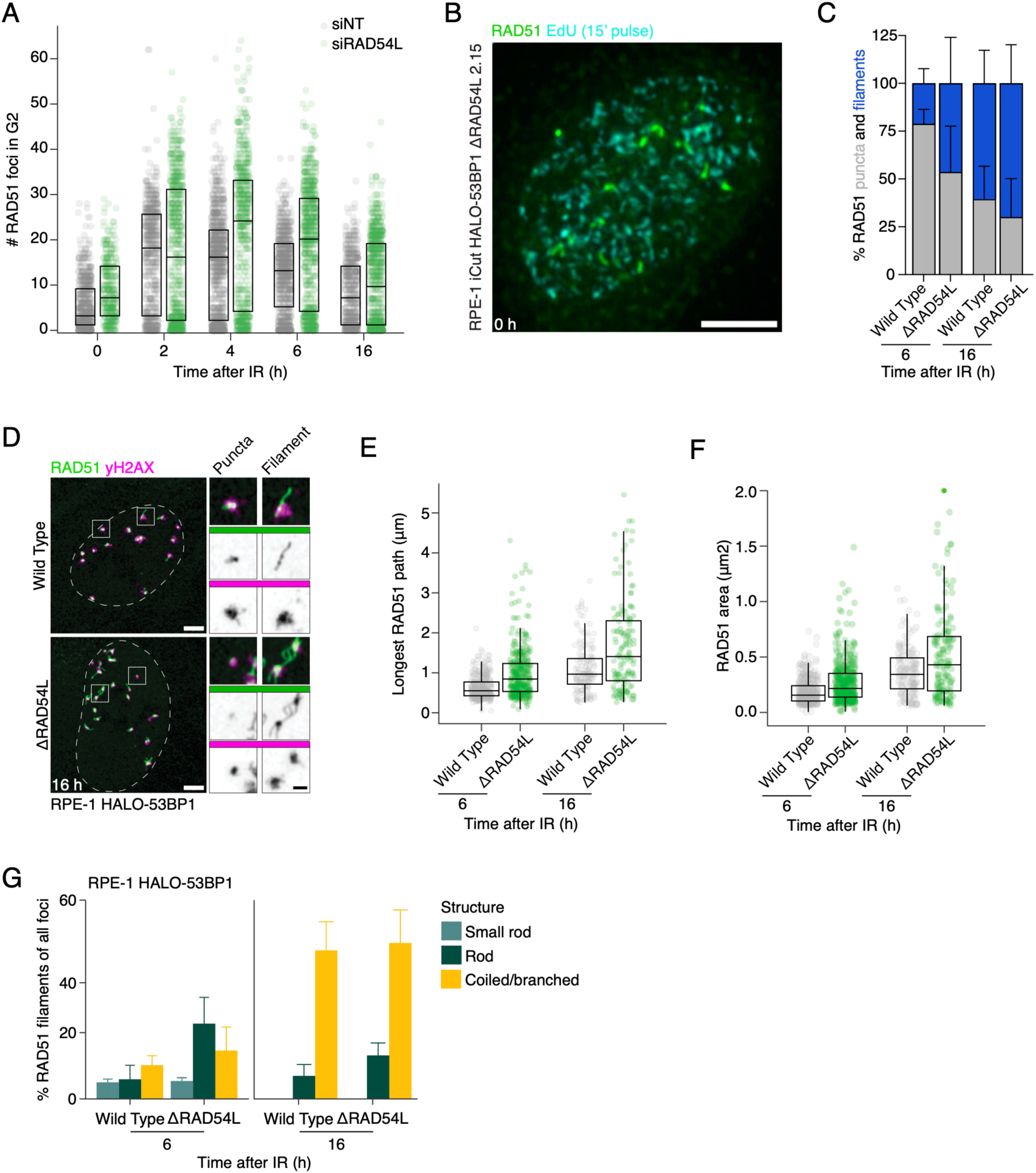
RAD54L is required for the resolution of RAD51 filaments. A) Immunofluorescence staining of RAD51 foci in RPE-1 HALO-53BP1 treated with siNT and siRAD54L during G2 phase of the cell cycle at different time points after irradiation (4 Gy). Three independent replicates are displayed. B) Immunofluorescence staining of RAD51 (green) puncta and combined with EdU (cyan) in an RPE-1 HALO-53BP1 ΔRAD54L cell. EdU was added 15 min before fixation to resolve filament structures in S-phase cells by super-resolution. Scale bar 5 μm. C) Average percentages of RAD51 filaments in RPE-1 HALO-53BP1 wild-type and ΔRAD54L cells at 6, 16 hours after irradiation (4 Gy). Data were obtained through Ilastik. D) Super-resolution microscopy of RPE-1 HALO-53BP1 wild-type and ΔRAD54L cells exposed to irradiation (4 Gy,16 hours), immunostained with yH2AX (magenta) and RAD51 (green) showing both puncta and filaments. Scale bars, 4 μm. Insets 1 μm. E) Ilastik analysis of the area of all RAD51 foci RPE-1 HALO-53BP1 wild-type and ΔRAD54L cells at 6, 16 hours after irradiation (4 Gy). Data are presented with individual measurements and box plots. Centre line, median; box limits, 25th and 75th percentiles. F) Ilastik analysis of the longest RAD51 path of every individual RAD51 foci in RPE-1 HALO-53BP1 wild-type and ΔRAD54L cells at 6, 16 hours after irradiation (4 Gy). Data are presented with individual measurements and box plots. Centre line, median; box limits, 25th and 75th percentiles. G) Average percentages of different RAD51 filament assemblies at 6 and 16 hours after irradiation (4 Gy) in RPE-1 HALO-53BP1 wild-type and ΔRAD54L cells. Small rods (light green), rods (dark green) and coiled/branched (yellow) were used as categories. Data were obtained through Ilastik. N=2.

### Visual evidence of independent versus coordinated search modes

Our data thus far show that RAD51 filaments grow in size over time and adopt altered, more complex, topographical structures (coiled/branched) (Fig.2A,D). This growth is further enhanced if strand invasion is inhibited by loss of RAD54L function. Branched filaments often adopted I-, Y- or V-shapes (Fig 4A,B). We hypothesized that these structures could represent different modes of homology search. In the case of Y or V-shapes, the two resected DNA ends from either side of the break would seek homology independently, whereas I-shapes would represent the coordinated search of these ends (Fig 4A,B). Branched structures could very commonly be observed at a later time point, in both the wild-type controls and the RPE- 1ΔRAD54L cells (Fig. 3G, 4C). To find further support for the notion that Y and V-shapes might represent DNA ends engaged in independent search modes, we analyzed the orientation of the Y- and V-shapes to several known DNA damage factors, such as yH2AX, RPA2 and BRCA2 (Fig 4D, Suppl. Fig.4A,B). The yH2AX core is commonly associated with the brightest part of the Y- or V-shape, consistent with a model in which the probing ends split in an independent search mode (Suppl. Fig.4A). RPA2 likewise decorates this DNA damage core in an ordered, circular arrangement in both wild-type as well as ΔRAD54L cells (Fig.4D). BRCA2 by contrast coats the entire filament (Suppl. Fig.4B). Thus, RAD51 filaments extend out from the core where RPA2 is assembled, sometimes with both ends paired (I-shape) and sometimes with both ends split (Y- or V-shapes), possibly representing alternate modes of homology search.

**Fig 4:**
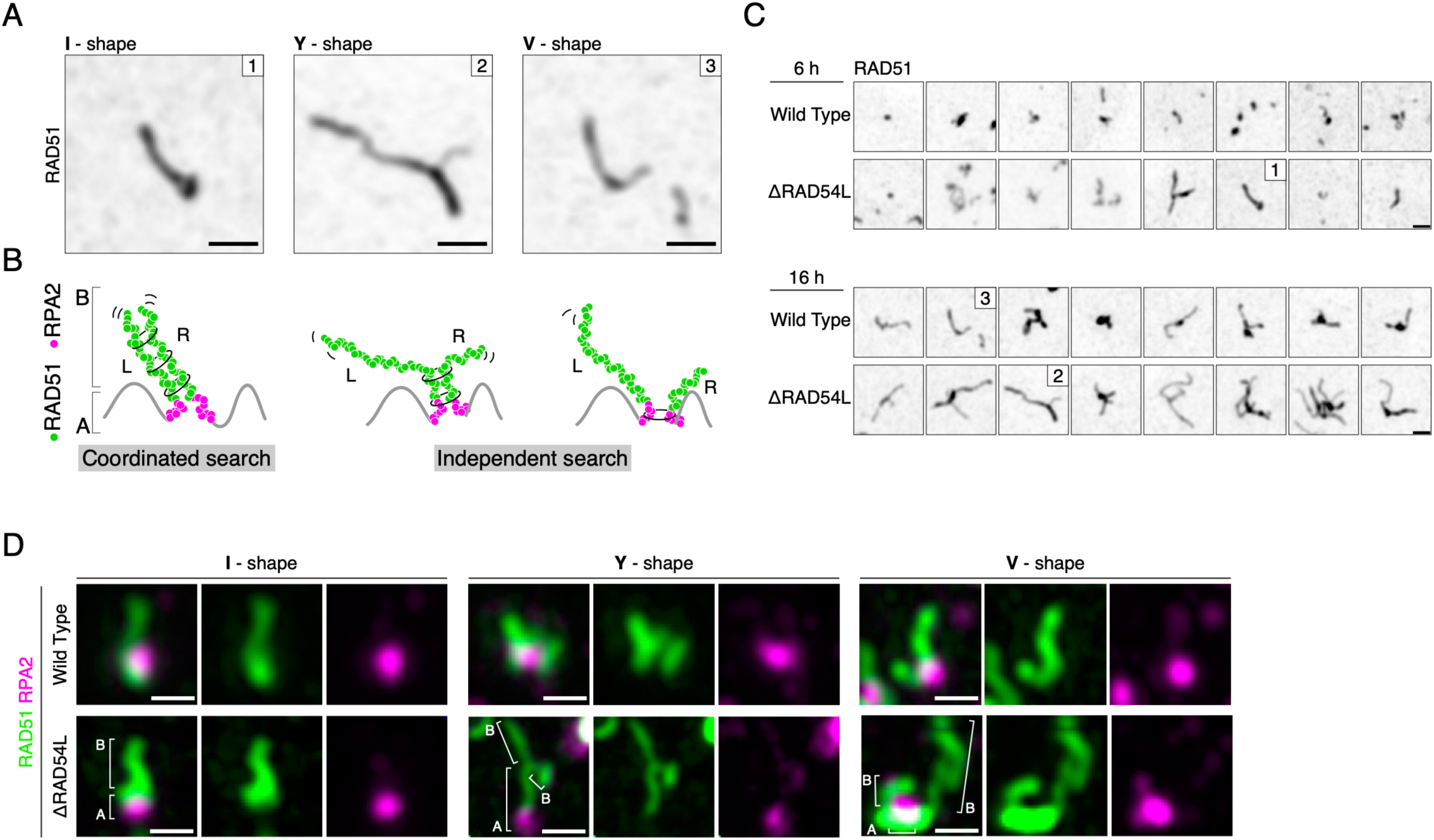
Visual evidence of independent versus coordinated search modes. A) Magnifications of the three images from (C) depicting examples of 1) I-shape, 2) Y-shape or 3) V-shape RAD51 filaments. Underneath the respective images Where the broken DNA seeks for its donor via dependent (1) or/and independent (2+3) homology search mechanisms. B) Different models for homology search. C) Super-resolution images of RPE-1 HALO-53BP1 wild-type and ΔRAD54L cells exposed to irradiation (4 Gy) for the indicated times, immunostained with RAD51. Scale bars, 0.5 μm. Insets, 1 μm. D) Super resolution of immunostained RAD51 (Green) and RPA2 (Magenta) in RPE-1 HALO-53BP1 wild-type and ΔRAD54L cells (16 hours, 4 Gy). Scale bar, 0.5 μm.

### RAD51 filaments recruit MND1

To obtain more insights into the real-time dynamics of RAD51 nucleoprotein filaments in human cells, we tried to produce a fluorescently tagged variant of RAD51. Due to technical difficulties however, our attempts unfortunately failed. We therefore took an alternative approach, based on our recent observations that MND1 acts as a cofactor in RAD51-mediated repair in human somatic cells and its dynamic localization can be traced in living human cells using GFP-MND1 ^31^. Importantly, when examining RAD51 foci observed at 6 and 16 hr after irradiation, we find that GFP-MND1 colocalizes with RAD51 in damage-induced foci (Fig.5A), with a near-complete overlap in staining pattern at 16 hours post-damage (Fig.5B). In addition, GFP-MND1 is also present on the RAD51 filaments (Fig.5C), that can be observed in I, Y, or V-shaped structures (Fig.5C), and that grow longer and become less spherical at later time points after irradiation (Suppl. Fig. 5A). GFP-MND1 filaments form on the core of a 53BP1 focus, and this core stains positive for γH2AX (Suppl. Fig.5B), BRCA1 and BRCA2 (Suppl. Fig.5C). The GFP-MND1 filaments display a similar variety in topographical structures as we observed for RAD51 filaments (Suppl. Fig.5B, D). 2D-STORM imaging of RPE-1 cells expressing GFP-MND1 revealed a clear filamentous appearance of MND1-positive foci at 16 hours post-damage (Fig.5D). Taken together, these data demonstrate that GFP-MND1 consistently co-localizes to the RAD51 nucleoprotein filaments and can be used to track the dynamics of these filaments.

**Fig 5:**
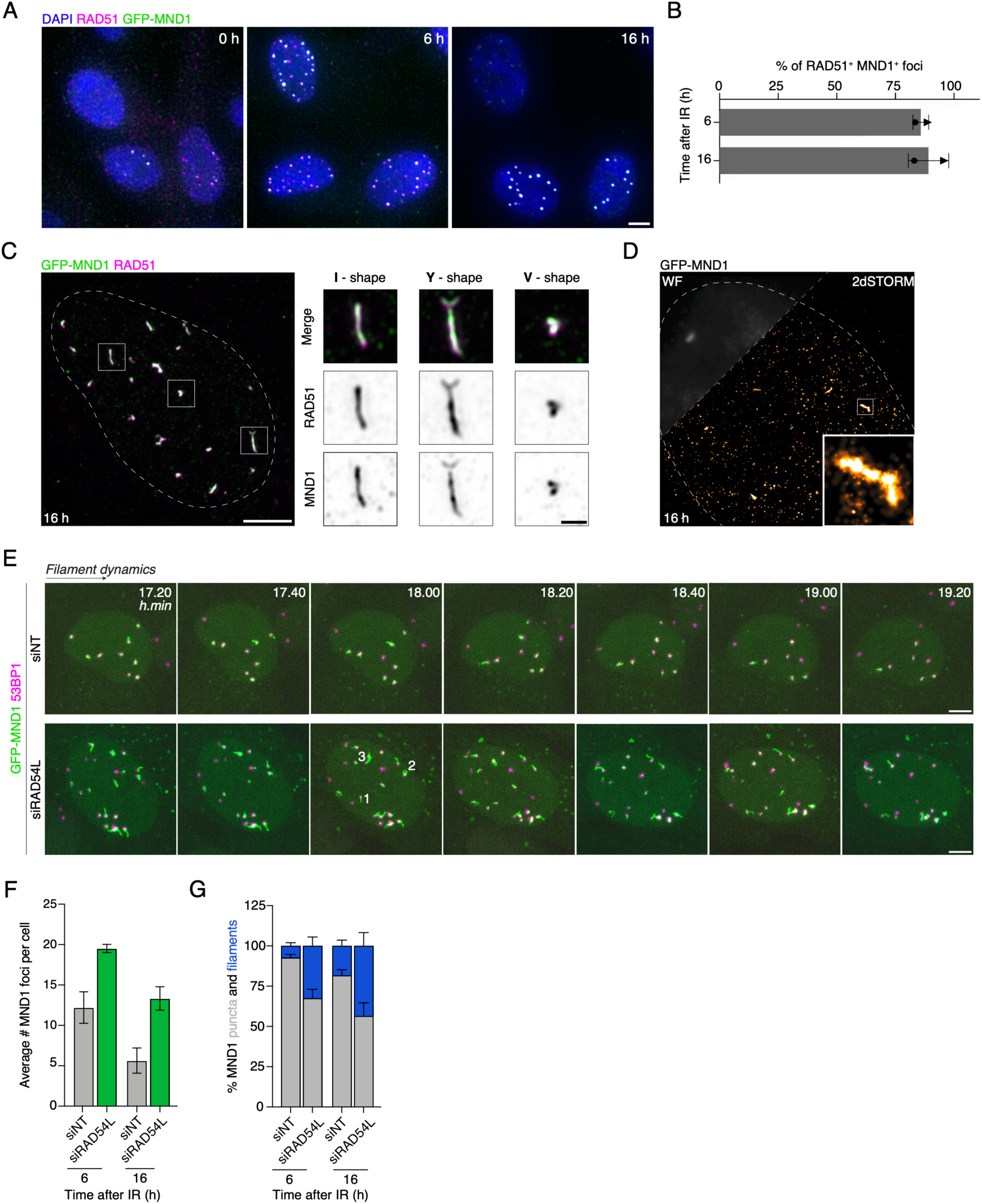
RAD51 filaments recruit MND1. A) Examples of wide-field images of wild-type RPE-1 HALO-53BP1 GFP-MND1 cells exposed to irradiation (4 Gy) for the indicated times. Cells were immunostained with RAD51 (magenta), GFP (green) and DNA (DAPI). Scale bars, 5 μm. B) Quantifications of the average percentages RAD51+ foci that are also MND1+ that 6 and 16 hours post damage. N=2 (>250 cells per replicate). C) Super-resolution of immunostained GFP-MND1 (green) and RAD51 (magenta) RPE-1 HALO-53BP1 GFP-MND1 wild-type cells showing overlap of the proteins. Scale bars, 5 μm. Magnifications of the three examples where for MND1 the I-shape, V-shape and V-shaped RAD51 (magenta) / MND1 (green) filament is observed. Insets, 1 μm. D) Wide-field (WF, gray) and 2DSTORM image of the same wild-type RPE-1 HALO-53BP1 GFP-MND1 immunostained with GFP to visualize a continuous MND1 filament. 16 hours post 4 Gy. E) Stills of a live-cell imaging movie of RPE-1 HALO-53BP1 (magenta) RPA1-mScarlet (not shown) GFP-MND1 (green) cells exposed to irradiation (4 Gy) for the indicated times. Cells were treated for 48h with a siRNA NT or RAD54L before irradiation. Scale bars, 5 μm. F) Average percentages of MND1 foci 6 and 16 hours post damage. Quantifications from live-cell movie. N=2. G) Average percentages of MND1 filaments at 6, 16 hours post damage. Quantifications from live-cell movie. N=2.

To investigate if filament formation is a peculiar feature of GFP-tagged MND1, we obtained an RPE-1 cell line expressing exogenous MND1 with a HALO-tag (Suppl. Fig.5E). This tagging strategy has the added benefit that it is compatible with the FUCCI system ^32^ (Suppl. Fig.5E) allowing for real-time cell cycle stratification during filament formation. Using HALO-MND1-RPE1-FUCCI cells we could show that HALO-MND1 filaments are formed in S and G2 phase cells at 6 to 16 hr after irradiation (Suppl. Fig. 5F-H), very similar to what we have seen for RAD51.

Next, we asked if the GFP-MND1 filaments respond to the depletion of RAD54L. In RAD54L-depleted cells, GFP-MND1 filaments grew larger (Fig.5E), and the overall number of MND1 foci that was induced by irradiation was increased (Fig.5E,F). Moreover, the resolution of filaments between 6 and 16 hr appears to be inhibited (Fig.5E) and the relative percentage of MND1-coated filaments was enhanced after depletion of RAD54L (Fig.5E,G), consistent with what we observed for RAD51.

### Real-time dynamics of MND1 filaments

To gather further evidence that RAD51/MND1-positive filaments represent homology search in action in living human cells, we monitored GFP-MND1 behavior after irradiation over longer periods of time using live-cell imaging. To track the core of the break in these experiments, we made use of a HALO-tagged version of 53BP1. To this end, we expressed GFP-MND1 in a RPE-1 cell line containing endogenously tagged HALO-53BP1. At 5-30 min after ionizing radiation (4 Gy), single repair foci appear, marked by loading of HALO-53BP1, consistent with previous reports ^33,34^. Time-lapse imaging showed that 53BP1-positive foci are initially devoid of GFP-MND1, but recruit GFP-MND1 to a focal site somewhere on the periphery of the 53BP1 focus (Fig.6A). These foci extend to filaments (Fig.6B), some of which are eventually resolved, shortly followed after by resolution of the 53BP1 focus (Fig.6B). GFP-MND1 filaments appear to extend away from the core and display a constant process of stretching and retracting over time (Fig.6C), potentially to locate the donor template. Live cell imaging showed that the filaments can span very large distances. This, combined with movements in different directions away from the core of the focus could allow the nucleoprotein filaments to explore very large volumes in the nucleus. This highly dynamic nature of MND1 mirrors the characteristics of Rad51 filaments observed in yeast ^22^ and is the first live visualization of homology search in mammalian cells.

**Fig 6:**
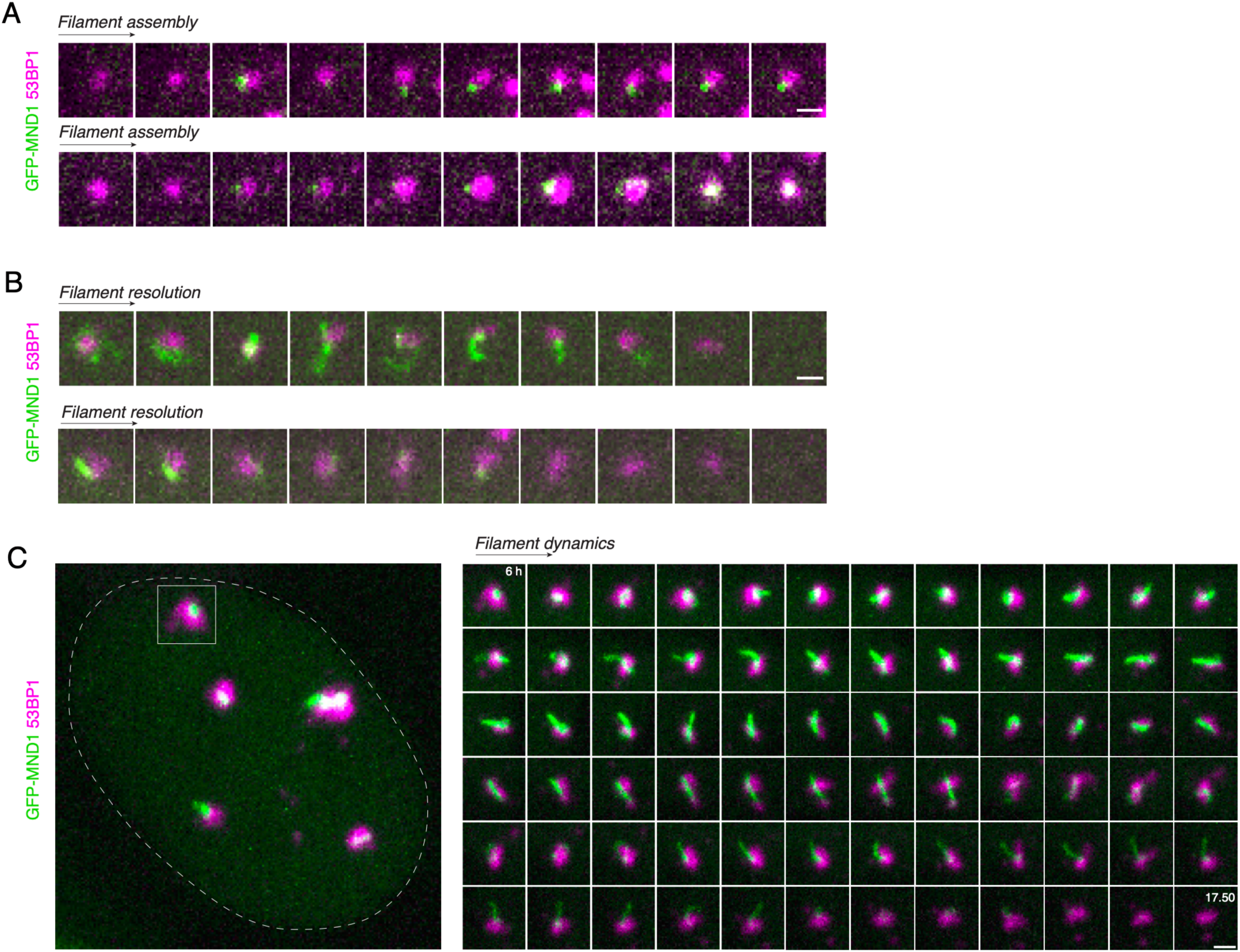
Real-time dynamics of MND1 filaments. A) Stills of a live-cell imaging movie showing the assembly of a MND1 (green) filament from a 53BP1 focus in a wild-type RPE-1 HALO-53BP1 (magenta) RPA1-mScarlet (not shown) GFP-MND1 (green) cell exposed to irradiation (4 Gy) for the indicated time. Time interval between images is 10min. Scale bars, 0.5 μm. B) Stills of a live-cell imaging movie showing the resolution of a MND1 (green) filament into a disappearing 53BP1 focus in a wild-type RPE-1 HALO-53BP1 (magenta) RPA1-mScarlet (not shown) GFP-MND1 (green) cell exposed to irradiation (4 Gy) for the indicated time. The time interval between images is 10min. Scale bars, 0.5 μm. C) Stills of a live-cell imaging movie showing the assembly, dynamic movement and resolution of an MND1 (green) filament from and into a disappearing 53BP1 focus in a wild-type RPE-1 HALO-53BP1 (magenta) RPA1-mScarlet (not shown) GFP-MND1 (green) cell exposed to irradiation (4 Gy) for the indicated time. The time interval between images is 10min. Scale bars, 0.5 μm.

During these live cell imaging experiments, we observed the formation of I-shaped and V-shaped GFP-MND1 filaments that grew out from the 53BP1 core (Fig.7A). We also occasionally observed GFP-MND1 filaments that appear to connect multiple 53BP1-positive foci (Fig.7A). To investigate these bridging events in more detail, we performed time-lapse imaging in RPE-1 cells expressing HALO-53BP1, GFP-MND1 and RPA1-mScarlet, the latter protein to mark ssDNA. Using these cells we could show that both ends of the GFP-filaments that connect two 53BP1 foci are also positive for RPA1-mScarlet (Fig.7B) suggesting there is ssDNA on either side of these bridging filaments. Further analysis in fixed cells showed that bridging filament can also be observed between two RPA2-positive, γH2AX-positive, or BRCA2-positive regions (Suppl. Fig.6A). Important to note that while there invariably was a RPA2 focus associated with one end of the filament, the other connected RPA2 focus could also be observed along the length of the filament, rather than at the end (Fig.7C). This implies that these filaments have retained patches of RPA that have not (yet) been replaced by RAD51, or that RPA is recruited to the ssDNA in the D-loop that forms when the RAD51/MND1 nucleoprotein filament invades the dsDNA template (Suppl. Fig.7B-D).

**Fig 7:**
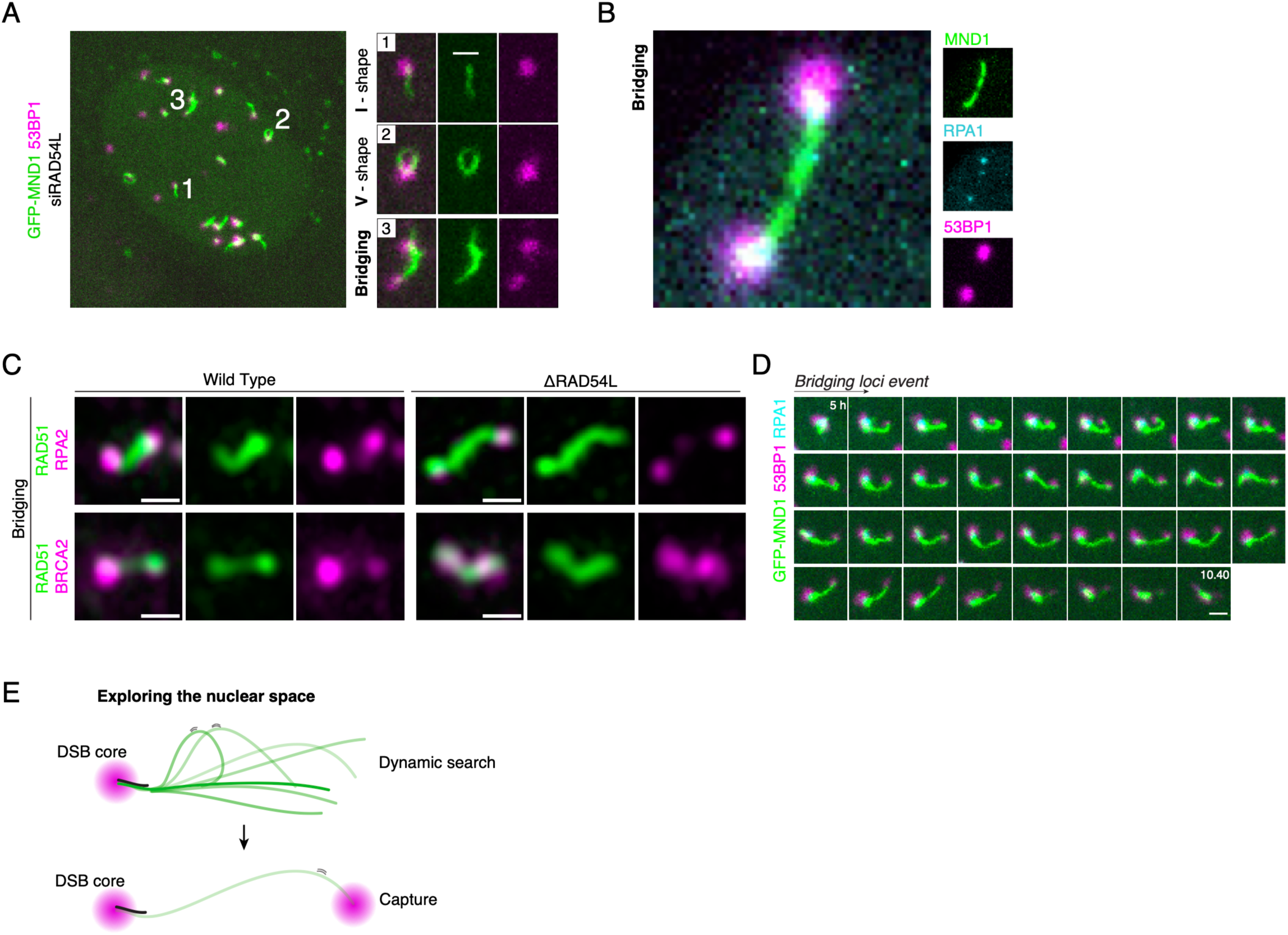
Real-time dynamics of MND1 filaments. A) Magnifications of the three stills from the live-cell movie from (A) depicting examples of 1) I-shape, 2) V-shape or 3) bridging RAD51 (green) filaments with 53BP1 foci (magenta). Insets, 1 μm. B) Magnification of a still from a live-cell movie of wild-type RPE-1 HALO-53BP1 (magenta) RPA1-mScarlet (cyan) GFP-MND1 (green) cells depicting a bridging MND1 filament between two 53BP1 and RPA1 foci. C) Super-resolution of immunostained RAD51 (green) and RPA2 (magenta) or RAD51 (green) and BRCA2 (magenta) in RPE-1 HALO-53BP1 wild-type and ΔRAD54L cells (16 hours, 4 Gy) highlighting the bridging event of the filament. Scale bar, 0.5 μm. D) Stills of a live-cell imaging movie following a wild-type RPE-1 HALO-53BP1 (magenta) RPA1-mScarlet (cyan) GFP-MND1 (green) filament exposed to irradiation (4 Gy) for the indicated times. Scale bars, 0.5 μm. E) Model: A RAD51 filament (green) forms from the ‘core’ of the DSB (magenta). It dynamically explores the nuclear environment, navigating through the crowded 3D nucleus to seek its homologous sequence. Upon capturing a sequence that appears homologous, a capture event occurs. If the sequence is heterologous, the filament releases it and continues searching. If the sequence is homologous, the repair process is continued.

When following connecting filaments in time lapse imaging experiments, we could observe examples where an originally “one-sided” filament gave rise to a second, less prominent 53BP1 focus that was brought in proximity to the original focus as the filament retracted (Fig.7D), suggestive of a search and capture event in HR (Fig. 7E).

### Cohesin, not CTCF, modulates RAD51-mediated repair

Our data thus far suggest that RAD51/MND1 filaments facilitate pairing between distant sequences, possibly leading to homology-directed strand invasion. Cohesin facilitates physical association and alignment of sister chromatids from replication onwards and loss of cohesin subunits has been shown to impair HR in various organisms ^35,36^. To test if cohesin controls the behavior of RAD51/MND1 filaments, we used a HCT116 cell line that expresses the essential cohesin subunit RAD21 endogenously tagged with a degron ^37^. This allows for rapid degradation of RAD21 upon the addition of 5-Ph-IAA (Suppl. Fig.7A). To test repair kinetics after cohesin depletion, 5-Ph-IAA was added 1 hour before irradiation to allow for complete RAD21 degradation, rendering the cohesin complex non-functional, as evidenced by a clear mitotic cohesin-defect (Suppl. Fig.7B). Therefore, we refer to the RAD21 depletion as cohesin depletion. Cohesin deficiency did not affect the global induction of damage as measured by the initial appearance γH2AX (Fig.8A,B), but the resolution of γH2AX foci at later stages was partially impaired (Fig.8A,B). Cohesin loss led to a more prominent impairment of the resolution of RAD51 foci (Fig.8A,C). Next, we examined the effects of cohesin loss on RAD51 filament topography using super-resolution imaging. While a significant fraction of the RAD51 filaments appears to have been resolved in HCT116 cells between 6 and 16 hr (Fig.8D), filaments continued to accumulate with more complex topographies during this time window in cohesin-depleted cells (Fig.8D). At 6 hours post-damage, RAD51 focus sizes were indistinguishable from control conditions, but at 16 hours, we more frequently observed elongated structures in cells that lacked cohesin (Fig.8E,F). This indicates that cohesin loss influences RAD51 dynamics and filament resolution in human cells, implying that cohesin loss should impair DNA damage repair in S/G2. Consistent with this notion, we find that loss of cohesin has a major effect on mitotic entry after DNA damage (Suppl. Fig.7C,D). Similarly, resolution of GFP-MND1 foci/filaments was impaired upon loss of cohesin (Suppl. Fig.7F-G). These data demonstrate that loss of cohesin impairs HR repair and perturbs mitotic progression under damaged conditions.

**Fig 8:**
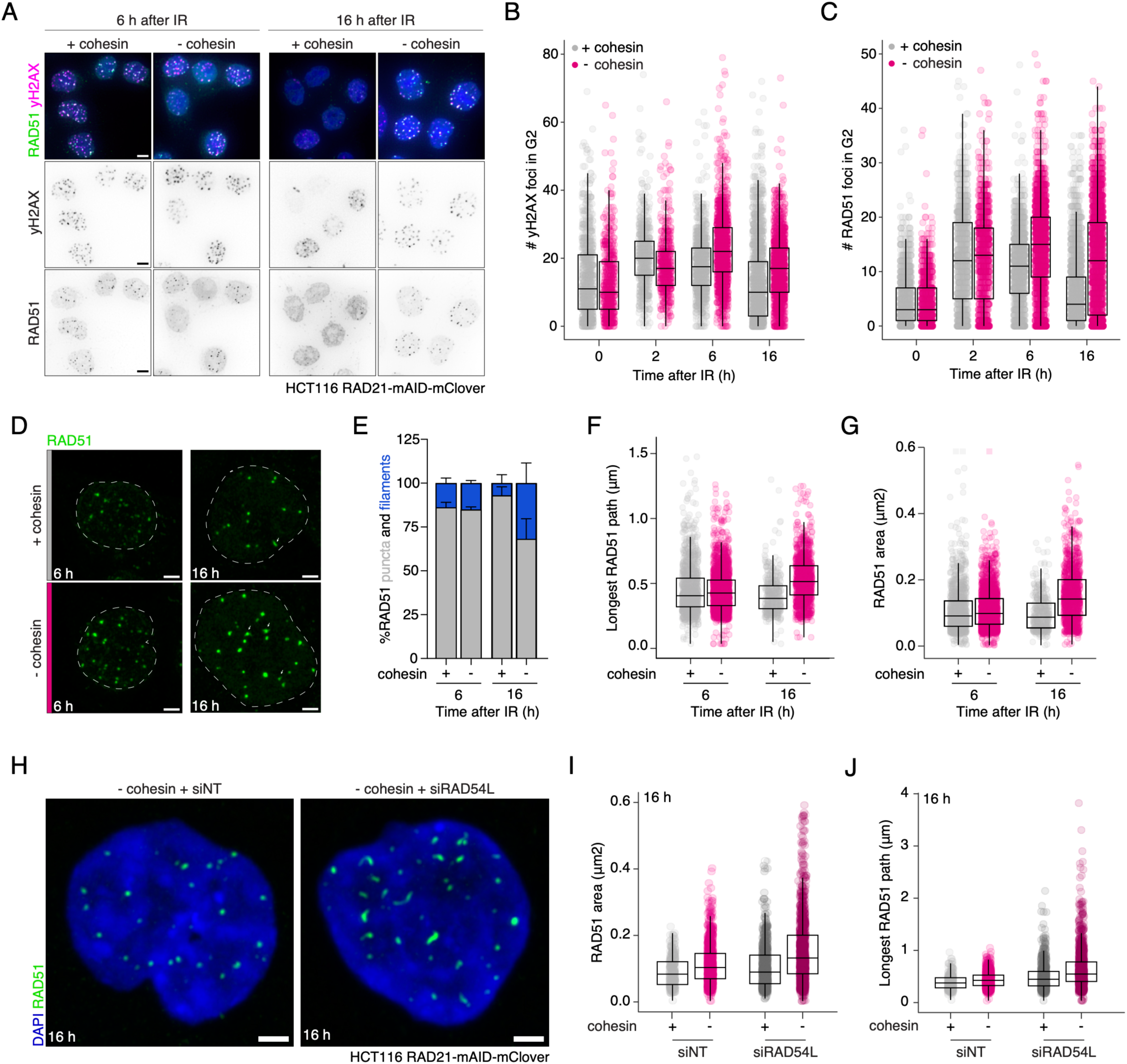
Cohesin and RAD54L are required for distinct steps of the homology search. A) Wide-field microscopy examples of HCT116 RAD21-mAID-mClover cells exposed to irradiation (4 Gy, 6 hours and 16 hours), immunostained with yH2AX (magenta) and RAD51 (green). Scale bars, 5 μm. Cells were either treated with DMSO (+cohesin) or 5-Ph-IAA to deplete RAD21 (-cohesin) 1 hour before irradiation. B-C) Quantification of the amount of yH2AX (B) or RAD51 (C) followed indicated hours after irradiation (4 Gy) as shown in (A). Box plots: centre line, median; box limits, 25th and 75th percentiles. N=3. D) Super-resolution images of RAD51 foci in HCT116 RAD21-mAID-mClover cells exposed to irradiation (4 Gy, 6 and 16 hours). Scale bars, 2 μm. E) Average percentages of RAD51 filaments at 6, 16 hours after irradiation (4 Gy). Data were obtained through Ilastik. N=2. F) Ilastik analysis of the area of all RAD51 foci in cells treated as in (D). Data are presented with individual measurements and box plots. Centre line, median; box limits, 25th and 75th percentiles. G) Ilastik analysis of the longest RAD51 path of every individual RAD51 foci in cells treated as in (D). Data are presented with individual measurements and box plots. Centre line, median; box limits, 25th and 75th percentiles. H) Super-resolution images of RAD51 foci in HCT116 RAD21-mAID-mClover cells exposed to irradiation (4 Gy, 6 and 16 hours). Cells were treated for 48h with siNT or siRAD54L followed by 5-Ph-IAA to deplete RAD21 (-cohesin) or DMSO (+cohesin) addition 1 hour before irradiation (4 Gy, 16 hours). Scale bars, 2 μm. I) Ilastik analysis of the area of all RAD51 foci in cells treated as in (E). Data are presented with individual measurements and box plots. Centre line, median; box limits, 25th and 75th centiles. N=2. J) Ilastik analysis of the longest RAD51 path of every individual RAD51 foci in cells treated as in (H). Centre line, median; box limits, 25th and 75th percentiles. N=2.

In addition to linking sister chromatids, cohesin also forms DNA loops on single chromatids, a role which has also been suggested to contribute to DNA repair by defining topologically associated domains (TADs) ^38^, which are flanked by CTCF sites. TADs restrict the spreading of γH2AX and aid in DNA repair ^39^. To test if the repair defects observed after cohesin loss are mediated by CTCF, we depleted CTCF 1 hour before irradiation in HCT116 CTCF-mAID-Clover cells ^40^ and observed efficient CTCF depletion (Suppl. Fig.8A). While depletion of cohesin significantly affected RAD51 foci resolution, CTCF depletion had no impact on RAD51 foci resolution and RAD51 filament length at 16 hr post-damage (Suppl. Fig.8B,C). Also, after low-dose irradiation (2 Gy) no difference in mitotic entry was observed when comparing CTCF-depleted HCT116 cells with unperturbed controls (Suppl. Fig.8D). Overall, these findings indicate that the essential role of cohesin in HR is independent of CTCF-anchored DNA looping.

### Cohesin and RAD54L are required for distinct steps of the homology search

To investigate if RAD54L and cohesin function at the same step during homology search, we performed siRNA-mediated knockdown of RAD54L in HCT116-RAD21-mAID-Clover cells. Super-resolution imaging showed that in RAD54L/RAD21 co-depleted HCT116 cells, RAD51 filaments where much more pronounced than after cohesin or RAD54L depletion alone (Fig. 8H, Suppl. Fig. 9A,B). Co-depletion of cohesin and RAD54L enhanced filament length, area as well as the amount of RAD51 foci (Fig. 8I,J, Suppl. Fig. 9C,D) compared to their single depletions. In this double depletion, the total area of RAD51 filaments was 0.14 μm2 and the filaments were 0.5 μm long. This implies that homology sampling by the RAD51 filament is impaired at multiple independent levels in the absence of RAD54L and cohesin. Thus, RAD54L/cohesin activities are needed for homology search, and the two have additive activities in this pathway.

## Discussion

In this study, we use quantitative high-content super-resolution microscopy to visualize DNA repair proteins active in homologous recombination. We uncover a previously unreported phenomenon in living human cells: the formation of dynamic filamentous structures at DNA break sites. We can visualize the formation of RAD51 filaments in fixed, and MND1 filaments in live cells. We propose that these filaments represent an intermediate state of homologous recombination of DNA breaks that search for homologous DNA sequences. Such a search mechanism has recently also been observed in yeast, suggesting a shared principle for the search for donor templates by DNA damage filaments.

We show that RAD51/MND1 filaments originate from a confined DNA damage core, stained by γH2AX, RPA1, RPA2, and 53BP1. These filaments arise after induction of DNA damage, and become prominent at 6 hours post-damage. We used live-cell imaging of GFP-MND1 to address the movement of filaments with respect to the DNA damage core. We observe that GFP-MND1 filaments indeed originate and extend from this 53BP1-positive core, that these filaments adopt various conformations, and that the filament eventually retracts towards this core. The length of the RAD51 filaments aligns with the distance between sister chromatids observed previously ^25,26^. As the sister chromatid is often used as repair template, these findings together support a model in which extended filaments represent HR intermediates that seek for homology.

Homologous sequences of DNA breaks can also be located on homologous chromosomes. We observe that RAD51/MND1 filaments at 16 hours are enlarged. It is possible that the gradual extension of filaments represents the shift from local search (sister chromatid) towards long-range/global search (homologous chromosome). Indeed, recent experiments in yeast reveal that the search of a DNA break for donor sequences is restricted to its local environment shortly after break induction ^41^. An alternative explanation for the progressive extension of RAD51/MND1 filaments, is that breaks engage in HR, but that the subsequent localization of a donor template sequence fails. Excessive resection of these unrepairable breaks would then result in long stretches of ssDNA that are coated with RAD51/MND1. RAD51/MND1 filaments therefore may thus also represent products of failed repair. However, as these filaments sometimes resolve, presumably representing the repair of the break, we favour the model in which extended filaments represent HR intermediates that aid in the localization of repair donor templates.

It is likely that excessive resection of long stretches of ssDNA allows for the extension of RAD51 filaments. However, a filament could also extend when MND1 and/or RAD51 oligomerize independent of ssDNA binding, when the filament unfolds from a folded conformation into a more linearly-stretched structure ^42–44^, or when aligned RAD51 bundles (double-coated RAD51 filaments) shift towards lateral filaments (single-coated RAD51 filament) ^13,15,20^. This latter scenario could also explain the thicker central regions observed in the branched GFP-MND1 structures. It therefore will be interesting to test which of these activities control filament length.

Notably, the filaments often adopt I-, V- or Y-shaped conformations during this process. The I- conformation could represent a structure in which the two broken ends remain paired and move together, while V- and Y-shaped structures could represent the independent movement of two complete or partial loose ends respectively. Single-strand Hi-C studies in yeast ^41^ suggests that the two ends of a break often perform search for homologous sequences in a coordinated manner, but that they indeed not always remain together. We therefore propose that RAD51/MND1 filaments may search for homologous sequences outside the core of the DNA damage focus via coordinated and independent search mechanisms.

We sometimes observe that RAD51 filaments are flanked by RPA, 53BP1 and yH2AX. We speculate that this conformation reflects the capture event during homology search when RPA on one end binds ssDNA at the ssDNA/dsDNA transition of the break, and at the other end the ssDNA in the D-loop of the synaptic complex ^45,46^. In this model, RPA would then stabilize the ssDNA when homology is found by the RAD51 filament to prevent the formation of secondary structures ^47,48^. Another possibility is that the flanking RPA/53BP1 or yH2AX represent the junction between the ssDNA and dsDNA of both of the ends of the DSB. The RAD51-filaments in between these junctions are in the middle kept together via an end-tethering process (Figure 9). Maintenance of end-tethering involves several components during DSB repair in budding yeast Saccharomyces cerevisiae. This is initially mediated by the MRN complex and is followed by Exo1 and the Ddc1-Mec3-Rad17 checkpoint clamp (human 9-1-1 clamp) and cohesin independently ^49–52^. This tethering at the level of the resection could also be required for mammalian search mechanisms.

**Fig 9:**
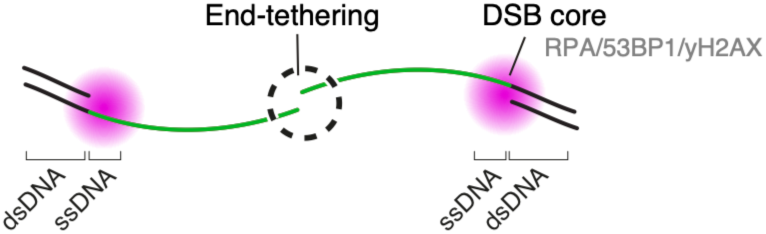
In this model the flanking RPA/53BP1 or yH2AX represent the junction between the ssDNA and dsDNA of both of the ends of the DSB. The RAD51-filaments in between these junctions are in the middle kept together via an end-tethering process.

We investigated the effect of loss of known homologous recombination factors on RAD51/MND1 filament dynamics. We focussed on RAD54L and cohesin, two factors that previously have been identified to control homologous recombination repair. RAD54L, a member of the SWI2/SNF2 family of DNA-dependent ATPases, facilitates strand invasion and promotes the dissociation of RAD51 from the D-loop or heterologous sequences. Cohesin by contrast builds loops to promote the local search for homology, facilitates the formation of γH2AX domains, and confers sister chromatid cohesion. We reveal that loss of either of these factors results in extended filaments compared to wild-type cells. Future studies are required to test which function(s) of RAD54L and cohesin restrict filament formation.

The ability to track the dynamics of homologous recombination in living cells provides valuable opportunities for screening genetic factors and small molecules that influence the homology search process. Through this approach, we demonstrate that filament length and resolution are controlled by RAD54L and cohesin. RAD54L likely guides the RAD51-coated ssDNA to its homologous sequence and displaces RAD51 from the D-loop. Cohesin, on the other hand, ensures optimal 3D chromatin organization to facilitate an efficient search-and-capture process. The transient and reversible RAD51/MND1 filaments visualized here for the first time in human cells serve as a powerful tool for understanding the dynamics of homology search and the impact of perturbations on genome stability in health and disease.

## Materials and Methods

### Cell culture and generation of CRISPR-Cas9 edited cell lines

RPE-1 cells were cultured in Dulbecco’s modified Eagle’s medium/Nutrient Mixture F- 12 (DMEM/F12, Gibco) supplemented with 6% fetal bovine serum (FBS, Sigma), 1% GlutaMAX (Gibco), and 100 U·mL^−1^ penicillin-streptomycin. HAP1 (Brummelkamp Laboratory, Amsterdam, The Netherlands) cells were cultured in Iscove’s modified Dulbecco’s medium (IMDM, Gibco) supplemented with 12% FBS, 1% GlutaMAX (GIBCO), and 100 U·mL^−1^ penicillin–streptomycin. U2OS were cultured in DMEM (GIBCO) supplemented with 6% FBS, 1% GlutaMAX (Gibco), and 100 U·mL^−1^ penicillin–streptomycin. HCT116 cells were cultured in RPMI 1640 (Gibco) supplemented with 12% FCS, 1% GlutaMAX (GIBCO), 1% HEPES (GIBCO), 1% MEM nonessential amino acids (GIBCO, Waltham, USA), and 100 U·mL^−1^ penicillin–streptomycin. All cell lines were tested negatively for mycoplasma contamination before experiments were performed.

For gene-editing experiments, 250.000 cells were plated in a six-well dish, the next day, cells were transfected a synthesized guide RNA (IDT) and the corresponding trans-activating CRISPR (tracr) RNA (IDT) that were mixed in equimolar concentrations to create a final duplex of 20 nM in OptiMEM (Gibco). The mixture was then transfected using RNAiMAX (Invitrogen) in Cas9 expressing RPE-1 cells (Dox/Shield-1 induced) or HAP1 pcW-Cas9 cells. Twenty-four hours later, the media was refreshed and cells were grown for an additional 48 hours. Single-cell clones were derived by limiting dilution. Successful editing of selected clones was verified by using TIDE analysis ^53^ using the Sanger sequencing of the polymerase chain reaction (PCR) products. Loss of the targeted protein was also assessed by immunoblotting.

### Western Blot Analysis

Western blot analysis was conducted as previously described ^31^. Briefly, proteins were separated by SDS-polyacrylamide gel electrophoresis and transferred onto nitrocellulose membranes. The membranes were then blocked in 5% BSA or 5%MILK in TBS-T and incubated with primary antibodies overnight at 4°C, with antibody dilutions specified below. Following this, secondary antibodies were applied for 1 hour at room temperature. Protein visualization was achieved using enhanced chemiluminescence (ECL; GE Healthcare, Chicago, USA). Antibodies used:

### Antibodies and chemicals

The following primary antibodies were used in this study: anti-H2AX ser139 (yH2AX, 05-636, Upstate, Burlington, USA, 1: 500 IF, 1:1000 WB), anti-RAD51 (ab63801, Abcam, Cambridge, UK, 1:500 IF, anti-GFP (11814460001, Roche, Basel, Switzerland, 1:1000 WB/IF), anti-HSP90 (sc7947, Santa Cruz, Dallas, USA, 1 : 1000 WB). The following secondary antibodies were used for western blot experiments: peroxidase-conjugated goat anti-rabbit (P448, DAKO, Santa Clara, USA, 1:1000) and goat anti-mouse (P0447, DAKO, Santa Clara, USA, 1:1000). Secondary antibodies used for immunofluorescence were goat anti-mouse-Alexa 488 (A11029, Mol Probes, Waltham, USA, 1:1000) and goat anti-rabbit-Alexa 568 (A11011, Mol Probes, Waltham, USA, 1:1000). Chemicals used in this study: neocarzinostatin (NCS, N9162, Sigma, Waltham, USA). Halo ligand Janilla fluor (Promega), 5-Ph-IAA (BioAcademia, Japan, #30-003).

### siRNA and crRNA Transfections

For siRNA transfections, RNAiMAX (Invitrogen, Waltham, USA) was used according to the manufacturer’s guidelines. The following siRNAs were employed in this study: non-targeting siNT and siRAD54L smart-pool all obtained from Dharmacon (Cambridge, UK). For crRNA transfections, the crRNAs were designed to target specific exons or sequences. These crRNAs were transfected using RNAiMAX in OptiMEM (GIBCO, Waltham, USA). The transfection procedure involved incubating 20 nM crRNA and tracrRNA with RNAiMAX for 20 minutes before adding the mixture to the cells. Indel generation in the RAD54L locus was confirmed using TIDE analysis of PCR products.

### Ionizing Radiation and Clonogenic Outgrowth

Cells were subjected to ionizing radiation using a Gammacell Exactor (Best Theratronics, Ottawa, Canada) equipped with a 137^C^s source. To assess the sensitivity of various cell lines to γ-irradiation, a low density of cells was plated per well. These cells were then treated with different doses of irradiation and allowed to grow into single colonies over a period of 6 days. After this incubation period, the number of colonies formed was counted and normalized to the number of colonies in the untreated control condition.

### Constructs

pCW_HALO-MND1 plasmid was cloned by PCR amplification of MND1 from a construct previously generated ^31^ using the following primers:

fwd 5′- AGCGCTTCGGCTGGAGGAAGTGGTtcaaagaaaaaaggactgagtgca-3′

rev 5′- AAAAGGCGCAACCCCAACCCCACGCGTttagtctatgtagtcaaagtcttctggaattcc-3′ and HALO from plasmid DNA with the following primers:

fwd1 5′- TCGCCTGGAGAATTGGatggcagaaatcggtactggc-3′

rev1 5′- CAGCCGAAGCGCTGCCGGAAATCTCGAGCGTAc-3′.

fwd2 5′-GCTCTACAAAAGCGCTTCGGCTGGAGGAAGTGGTtcaaagaaaaaaggactgagtgca-3′.

rev2 5′-AAAAGGCGCAACCCCAACCCCACGCGTttagtctatgtagtcaaagtcttctggaattcc-3′.

The PCR products were ligated into a cut-open (AgeI, EcoRI) pCW vector with BFP-T2A-Blast selection marker by Gibson ligation. Plasmids were then lentivirally transduced into RPE-1 and HCT116 cells and selected with blasticidin.

### Lentiviral transductions

HEK293T cells (5× 106) were plated in a 10-cm dish the day before they were transfected with packaging plasmids: 4μg of pMD2.G (Addgene,12260), 4 μg of psPAX-2 (Addgene,12259), and 4μg of the target vectors using PEI. Viral supernatant was collected 48 hours after transfection, filtered (0.45 μm syringe), and used for transduction in the presence of polybrene (8μg/ml; Sigma-Aldrich). To select out positive transduced cells, cells were treated with blasticidin (15μg/ml) until all non-transduced well died.

### Immunofluorescence

Cells were seeded on coverslips (Thermo Fisher Scientific) and treated as indicated in the figure legends. To stain for GFP (11 814 40 001, Roche), RAD51 (anti-RAD51, ab63801 abcam), or yH2AX (yH2AX, 05-636 Upstate), (RPA2, Calbiochem), (BRCA2, Calbiochem) foci in cells, cells were fixed and permeabilizated with 4% PFA and 0.5% Triton X-100 in PBS, slides were incubated with primary antibodies in IF buffer for 1 hour at room temperature. Cells were then washed three times with PBS-T and subsequently blocked in PBS supplemented with 0.1% Tween-20 (PBS-T) and 5% bovine serum albumin (BSA) for 30 min. Primary antibody incubation was performed over-night at 4 °C (antibodies and dilutions stated below). Coverslips are washed with BSA in PBS-T, and secondary antibody (Invitrogen) and DAPI (2.5 μg/ml; Sigma-Aldrich) incubation is performed at room temperature for 1 h. Then coverslips were washed with PBS. EdU was stained by incubation in EdU staining buffer (100 mM Tris– HCl pH 8.5, 1 mM CuSO4), with 100 mM ascorbic acid and AF-647 azide (Invitrogen, 1/1000) for 30 min at room temperature. Then, coverslips were washed again and mounted with Prolong Gold (Invitrogen,) and stored at 4 °C.

All fluorescent images taken with the THUNDER microscope were acquired in three dimensions with a z-step of 500 nm. For optimal visualization the images are displayed as maximum intensity projection of the individual z-sections. Images were acquired on a THUNDER Imager (Leica Microsystems), equipped with 63X/1.40 Oil Obj. HC PL APO objective and deep-cooled 4.2 MP sCMOS camera and processed with thundering. For higher resolution, images were acquired on a Zeiss LSM 980 with Airyscan2 and 63X/1.4-0.6 Oil objective in super-resolution (SR) mode. After acquisition images were processed with ZEN software (Zeiss), applying the same processing strength per experiment. Using Imaris (Oxford Instruments), RAD51 structures were segmented with the surface algorithm and filtered by sphericity and longest axis using the bounding box measurement. Quantification of DNA damage–induced foci was performed in Fiji using CLIJ2/x libraries ^54^ with an in-house designed ImageJ macro (https://github.com/Bioimaging-NKI/Foci-analyzer), as described ^31^. Briefly, nuclei are segmented in the DAPI channel using StarDist ^55^. Total DAPI intensity is used to separate G1 and G2 phase cells. DNA damage foci are detected by applying marker-controlled watershed on local maxima in the foci channels, after which foci intensities and numbers are quantified per cell.

### 2dSTORM imaging and sample preparation

Samples were prepared as detailed described before ^56^. In brief, RPE-1 HALO-53BP1 GFP-MND1 cells were seeded on an ultra-clean coverslip. As described before, immunofluorescence staining was performed where after blocking with 5% Bovine Serum Albumin (BSA; Serva, Heidelberg, Germany) for 1 hour, cells were stained as follows. Cells were incubated with mouse anti-GFP (11 814 460 001, Roche). Subsequently all the cells were incubated with a goat anti-mouse antibody (Alexa-488, Thermo Fisher Scientific) and DAPI to stain the DNA at a final concentration of 0.01 mg/ml for 1 hour. The coverslip was mounted in a holder (Chamlide CMB mounting ring CM-B25-1, Live Cell Instrument, Seoul, South Korea) filled with 500 μl of OxEA imaging buffer. The OxEA buffer contains (50 mM β-MercaptoEthylamine hydrochloride (MEA, Sigma-Aldrich), 3% (v/v) OxyFlour™ (Oxyrase Inc., Mansfield, Ohio, U.S.A.), 20% (v/v) of sodium DL-lactate solution (L1375, Sigma-Aldrich) in PBS, pH adjusted to 8–8.5 with NaOH) as described before ^57^. Samples were imaged with the 488 nm/150 mW laser on a Leica SR-GSD microscope (Leica Microsystems, Wetzlar, Germany) equipped with an EMCCD camera (Ixon DU-897, Andor). We used a 160x oil immersion dedicated SR objective. Around 4000 frames were collected at 100 Hz with image size of 180×180 pixels. The data was analyzed with the ImageJ ThunderSTORM plugin ^58^, after background-correcting with a running-median background subtraction ^59^ using an in-house developed ImageJ plugin (https://github.com/Jalink-lab/Temporal-Median-Background-Subtraction).

### Pipeline for high-content analysis of filaments Quantification of RAD51 object Area and length

To quantify RAD51 object area and length, RAD51 structures were measured with Ilastik (v.1.3.2), using a pixel and an object classification. First, several FIJI in-house designed macros (https://github.com/Bioimaging-NKI/Visualising-homology-search-in-human-cells) were used to crop individual nuclei using the DAPI channel and semi-automatically segment RAD51 objects in 3D. The obtained label images were overlayed with raw data, allowing adjusting segmentation parameters. Label images were maximum intensity z-projected and used as input for Ilastik, where Object features (ObjectArea and diameter) were used for classification.

### Quantification of RAD51 structure

Using object classification in Ilastik the percentages of different RAD51 structures were quantified. First, representative images were used for manual annotation and training. For RAD51 structure quantification, the same FIJI macros were used before and 3D filaments stacks (50 × 50 pixels) were input and segmentation maps, generated with ImageJ were used as input. Size in pixels, mean intensity and total intensity were used for classification and output.

### Live-cell microscopy

For live-cellmicroscopy, RPE-1 HALO-53BP1 GFP-MND1 cells were seeded in a Lab-Tek II chambere coverglass (Thermofisher, 8-well) and kept overnight. Live-cell imaging was performed on a Andor Dragonfly spinning-disk confocal microscope system equipped with an Andor Dragonfly system on an inverted Leica DMI8 microscope with an incubation chamber (OKOlab) for temperature (37C) and CO2 control (5%). Images were acquired with a Leica 63x/1.47 NA Oil immersion objective and a sCMOS Zyla 4.2 (Andor). Images were acquired every 5 minutes for a time-course of 24h in three dimensions with *z*-steps spaced out by 500 nm. The max-projections were processed in FIJI applying a maximum intensity z projection. It is important to note that the above-described measures to enable three color live-cell imaging come at the expense of detailed z-depth.

## Supplemental figure legends

**Supplemental figure 1: RAD51 forms extended filaments at sites of DNA damage**

A) The whole image of the crops is shown in figure 1A. Immunofluorescence staining of RAD51 (green) puncta and filaments and γH2AX (magenta) in RPE-1 wild-type cells resolved by super-resolution. Cells were irradiated with 4 Gy and fixed 6 and 16 hours later. Scale bar, 5 μm. B) Immunofluorescence staining of RAD51 (green) in HAP1, U2OS and HCT116 cells resolved by super-resolution as filament structure throughout the nuclear volume. Scale bar 2 μm. A max projection is shown. C) Immunofluorescence staining of RAD51 (green) puncta and combined with EdU (cyan) in RPE-1 wild-type cells. EdU was added 15min before fixation to resolve filament structures in S-phase cells by super-resolution. Cells were irradiated with 4 Gy and fixed 16 hours later.

**Supplemental figure 2: RAD51 forms extended filaments at sites of DNA damage**

Ilastik analysis of the longest RAD51 path (A) and RAD51 area (B) of every individual RAD51 foci in RPE-1 wild-type cells at 6, 16h after irradiation (4Gy). Data were obtained through Ilastik. Experiments were biologically replicated twice. Data are presented with individual measurements and box plots. Centre line, median; box limits, 25th and 75th centiles.

**Supplemental figure 3: RAD54L is required for the resolution of RAD51 filaments**

A) Western blot confirming the two independent HAP1 pCW-Cas9 ΔRAD54L clones. B) Clonogenic outgrowth of HAP1 wild-type and two monoclonal ΔRAD54L cell lines (green) in response to increasing doses of IR. Data presented as mean ± SEM. N=3. C) Western blot confirming the two independent RPE-1 HALO-53BP1 ΔRAD54L clones. D) Immunofluorescence staining of RAD51 (green) puncta and filaments and γH2AX (magenta) in HAP1 wild-type and ΔRAD54L cells resolved by super-resolution. Cells were irradiated with 2Gy and fixed 6 and 24 h later.

**Supplemental figure 4: Two homology search modes: coordinated and independent**

A) Super-resolution images of RPE-1 HALO-53BP1 wild-type cells. Cells were exposed to irradiation (4 Gy) and fixed 16 hours later and immunostained with RAD51 (green) and γH2AX (magenta). From left to right: I-shape, Y-shape or V-shape RAD51 filaments. Where only part of the filaments co-localizes with γH2AX. Scale bars, 0,5 μm. B) Super-resolution of immunostained RAD51 (green) and BRCA2 (magenta) in RPE-1 HALO-53BP1 wild-type and ΔRAD54L cells (16 hours, 4 Gy). Scale bar, 0.5 μm.

**Supplemental figure 5: RAD51 filaments recruit MND1**

A) Example plot of the sphericity against the longest axis of GFP-MND1 foci measured in 3D, 6 or 24h after irradiation (4 Gy) of one cell. Plot of the sphericity and longest axis of all GFP-MND1 foci measured in many cells. The higher the value for sphericity, the more spherical (puncta) a focus is, the lower the value, the more elongated the focus is. Data were obtained through Imaris. B) Representative image of an RPE-1 wild-type cells immunofluorescence stained for GFP-MND1 (green) puncta and filaments and γH2AX (magenta) resolved by super-resolution. Cells were irradiated with 4 Gy and fixed 16 hours later. Scale bar, 4 μm. C) Examples of wide-field images of wild-type RPE-1 HALO-53BP1 GFP-MND1 cells exposed to irradiation (4 Gy) and fixed 16 hours after. Cells were immunostained with GFP (green), BRCA1 or BRCA2 (cyan) and DNA (DAPI). D) Stills of a live-cell imaging movie showing the assembly of a MND1 (green) filament from a 53BP1 focus in a wild-type RPE-1 HALO-53BP1 (magenta) RPA1-mScarlet (cyan) GFP-MND1 (green) cell exposed to irradiation (4 Gy). Depicting different conformations. E) HALO-MND1 overexpression induced by doxycycline addition in RPE-1 wild-type cells and RPE-1 FUCCI cells. F) Example image of a G2-phase, RPE-1 FUCCI HALO-MND1 cell showing filaments 16 hours after irradiation (4 Gy). G) Field of view of a live-cell imaging movie of RPE-1 FUCCI HALO-MND1 cells three cell cycle stages, Cell 1 = G2 (green), Cell 2 -S-phase (yellow) and Cell 3 = G1 phase (red). Image is taken 10 min before irradiation. H) Stills of a live-cell imaging movie showing the formation if HALO-MND1 foci in S and G2 phase, but not in G1 cells exposed to irradiation (4 Gy).

**Supplemental figure 6: The capture event in homology search**

A) Super resolution images of RPE-1 HALO-53BP1 wild-type cells. Cells were exposed to irradiation (4 Gy) and fixed 16 hours later and immunostained with RAD51 (green) and γH2AX (magenta). A bridging event is shown. Scale bar, 0,5 μm. B) Super resolution of immunostained RAD51 (green) and RPA2 (magenta) in RPE-1 HALO-53BP1 wild-type, showing a connected RPA2 dot along the RAD51 filament (16 hours, 4 Gy). Scale bar, 0.5 μm. C) Super resolution of immunostained RAD51 (green) and RPA2 (magenta) in RPE-1 HALO-53BP1 wild-type and ΔRAD54L cells (16 hours, 4 Gy) highlighting the connected dot along the filament. Scale bar, 0.5 μm. D) 2 potential models for the connected dots along a RAD51/MND1 filament. A RAD51 filament (green) forms from the ‘core’ of the DSB, which is covered by RPA2 (magenta). 1) There are alternating patches of RAD51/RPA2, where not all the RPA2 is replaced for RAD51. (2) The broken end dynamically explores the nuclear environment, navigating through the crowded nucleus to seek its homologous sequence. Upon capturing a sequence that appears homologous, a capture event occurs. This homology sampling can occur along an intersegmental stretch of DNA, possibly multiple times. If the sequence is heterologous, the filament releases it and continues searching. If the sequence is homologous, consecutive repair processes ensue.

**Supplemental figure 7: Cohesin, not CTCF, controls RAD51-mediated repair**

A) Scheme of procedure used for experiments performed with the HCT116 RAD21-mAID-mClover cells. Cells were either treated with DMSO (+cohesin) or 5-Ph-IAA (-cohesin) 1 hour before irradiation. Cells were fixed for immunofluorescence or harvested for western blot 6 or 16 hours after irradiation. Western blot showing near complete degradation of RAD21 after treatment with 5-Ph-IAA (-cohesin) for 7h in HCT116 RAD21-mAID-mClover cells. Western blot samples were harvested 6 hours after irradiation (4 Gy). B) Live cell imaging images of asynchronously growing HCT116 RAD21-mAID-mClover cells with DNA visualized using sir-DNA. Cells were either treated with DMSO (+cohesin) or 5-Ph-IAA (-cohesin) 1 hour before irradiation. Cells were imaged every 5min and their mitotic entry was assessed. Cell depleted for RAD21 (-cohesin) showed cohesion loss in mitosis. Scale bar 2 μm. C) HCT116 RAD21-mAID-mClover cells treated with DMSO (+cohesin) and 5-Ph-IAA (-cohesin) were left unirradiated (C) or irradiated with 2Gy (D) and subsequent imaged for 24h. Relative mitotic entry of asynchronous cells is shown. N=2. E) Insets of images with examples of the four categories of RAD51 assemblies observed in HCT116 RAD21-mAID-mClover cells. Small rods (light green), rods (dark green) and branched (yellow) were used as categories. F) Average percentages of theRAD51 filament assemblies at 6 and 16 hours after irradiation (4 Gy). RAD51 assemblies were scored as shown in (E). Data were obtained through Ilastik. N=2. G) Stills of a live-cell imaging movie of HCT116 RAD21-mAID-mClover (green) HALO-MND1 (magenta) cells treated with 5-Ph-IAA to deplete RAD21 (-cohesin) or DMSO (+cohesin) 1 hour before irradiation (4 Gy, 6 and 16 h). Clear loss of RAD21 is visible in the cells treated with 5-Ph-IAA. Images are acquired on a confocal microscope. H) Stills of a live-cell imaging movie of HCT116 RAD21-mAID-mClover HALO-MND1 (grey). Cells treated with 5-Ph-IAA to deplete RAD21 (-cohesin) or DMSO (+cohesin) 1 hour before irradiation (4 Gy). Clear MND1 foci accumulation is visible. I) Quantifications from live-cell movie in (H). N=3.

**Supplemental figure 8: Cohesin, not CTCF, controls RAD51-mediated repair**

A) Western blot showing near complete degradation of CTCF after treatment with 5-Ph-IAA (-CTCF) for 7h in HCT116 CTCF-mAID-mClover cells. Western blot samples were harvested 6 hours after irradiation (4 Gy). B) Quantification of the amount of RAD51 foci 6 and 16 hours after irradiation (4 Gy) in HCT116 CTCF-mAID-mClover cells. CTCF was depleted with 5-Ph-IAA (-CTCF) or left untreated 1 hour prior to irradiation. C) Ilastik analysis of the longest RAD51 path of every individual RAD51 foci in HCT116 CTCF-mAID-mClover cells at 6 and 16 hours after irradiation (4 Gy). Data were obtained through Ilastik. N=2. Data are presented with individual measurements and box plots. Centre line, median; box limits, 25th and 75th percentiles. D) HCT116 CTCF-mAID-mClover cells treated with DMSO (+cohesin) and 5-Ph-IAA (-cohesin) were irradiated with 2Gy (H) and subsequently imaged for 24h. Relative mitotic entry of asynchronous cells is shown.

**Supplemental figure 9: Double act: RAD54L and cohesin unite for effective homology search**

A) Super resolution images of HCT116 RAD21-mAID-mClover cells treated with siNT or siRAD54L for 48h. Cells were exposed to irradiation (4 Gy) for the indicated times, and immunostained with RAD51. Scale bars, 1 μm. B) Super resolution images of RAD51 foci in HCT116 RAD21-mAID-mClover cells exposed to irradiation (4 Gy). Cells were treated for 48h with siNT or siRAD54L followed by 5-Ph-IAA to deplete RAD21 (-cohesin) or DMSO (+cohesin) addition 1 hour before irradiation (4 Gy, 6 hours). Scale bars, 2 μm. C) Ilastik analysis of the area of all RAD51 foci in cells treated as in (C). Data are presented with individual measurements and box plots. Centre line, median; box limits, 25th and 75th percentiles. N=2. D) Ilastik analysis of the longest RAD51 path of every individual RAD51 foci in cells treated as in (C). Centre line, median; box limits, 25th and 75th percentiles. N=2.

## Acknowledgments

We kindly thank Masato T. Kanemaki for kindly sharing the HCT116 RAD21-mAID2-Clover and HCT116-CTCF-mAID2-Clover cells. We thank the members of the Medema Laboratory for helpful discussions. We thank the insightful discussion and comments of members from the Jacobs lab and the Rowland lab. We want to further thank Abdelghani Mazouzi, Gerben Vader and Amber Hondema for their useful input on this manuscript. This work was supported by funding from the Oncode Institute. The Oncode Institute is partly supported by KWF Dutch Cancer Society.

## Notes

https://github.com/Bioimaging-NKI/Visualising-homology-search-in-human-cells

### Competing Interest Statement

The authors have declared no competing interest.

## References

1. Li, X., and Heyer, W.D. (2008). Homologous recombination in DNA repair and DNA damage tolerance. Cell Res 18, 99–113. 10.1038/cr.2008.1.

2. Hu, J., and Crickard, J.B. (2024). All who wander are not lost: the search for homology during homologous recombination. Biochem Soc Trans 52, 367–377. 10.1042/BST20230705.

3. Bordelet, H., and Dubrana, K. (2019). Keep moving and stay in a good shape to find your homologous recombination partner. Curr Genet 65, 29–39. 10.1007/s00294-018-0873-1.

4. Renkawitz, J., Lademann, C.A., and Jentsch, S. (2014). Mechanisms and principles of homology search during recombination. Nat Rev Mol Cell Biol 15, 369–383. 10.1038/nrm3805.

5. Moynahan, M.E., and Jasin, M. (2010). Mitotic homologous recombination maintains genomic stability and suppresses tumorigenesis. Nat Rev Mol Cell Biol 11, 196–207. 10.1038/nrm2851.

6. Stark, J.M., and Jasin, M. (2003). Extensive loss of heterozygosity is suppressed during homologous repair of chromosomal breaks. Mol Cell Biol 23, 733–743. 10.1128/MCB.23.2.733-743.2003.

7. Kawabata, M., Kawabata, T., and Nishibori, M. (2005). Role of recA/RAD51 family proteins in mammals. Acta Med Okayama 59, 1–9. 10.18926/AMO/31987.

8. Sung, P., and Robberson, D.L. (1995). DNA strand exchange mediated by a RAD51-ssDNA nucleoprotein filament with polarity opposite to that of RecA. Cell 82, 453–461. 10.1016/0092-8674(95)90434-4.

9. Kong, M., and Greene, E.C. (2021). Mechanistic Insights From Single-Molecule Studies of Repair of Double Strand Breaks. Front Cell Dev Biol 9, 745311. 10.3389/fcell.2021.745311.

10. Cejka, P., and Symington, L.S. (2021). DNA End Resection: Mechanism and Control. Annual Review of Genetics 55, 285–307. 10.1146/annurev-genet-071719-020312.

11. Lehmann, C.P., Jimenez-Martin, A., Branzei, D., and Tercero, J.A. (2020). Prevention of unwanted recombination at damaged replication forks. Curr Genet 66, 1045–1051. 10.1007/s00294-020-01095-7.

12. Bell, J.C., Plank, J.L., Dombrowski, C.C., and Kowalczykowski, S.C. (2012). Direct imaging of RecA nucleation and growth on single molecules of SSB-coated ssDNA. Nature 491, 274–278. 10.1038/nature11598.

13. Chimthanawala, A., Parmar, J.J., Kumar, S., Iyer, K.S., Rao, M., and Badrinarayanan, A. (2022). SMC protein RecN drives RecA filament translocation for in vivo homology search. Proc Natl Acad Sci U S A 119, e2209304119. 10.1073/pnas.2209304119.

14. Wiktor, J., Gynna, A.H., Leroy, P., Larsson, J., Coceano, G., Testa, I., and Elf, J. (2021). RecA finds homologous DNA by reduced dimensionality search. Nature 597, 426–429. 10.1038/s41586-021-03877-6.

15. Lesterlin, C., Ball, G., Schermelleh, L., and Sherratt, D.J. (2014). RecA bundles mediate homology pairing between distant sisters during DNA break repair. Nature 506, 249–253. 10.1038/nature12868.

16. Wright, W.D., Shah, S.S., and Heyer, W.D. (2018). Homologous recombination and the repair of DNA double-strand breaks. Journal of Biological Chemistry 293, 10524–10535. 10.1074/jbc.TM118.000372.

17. Bonilla, B., Hengel, S.R., Grundy, M.K., and Bernstein, K.A. (2020). RAD51 Gene Family Structure and Function Annu Rev Genet. 54, 25–46. 10.1146/annurev-genet-021920-092410.RAD51.

18. Carver, A., and Zhang, X. (2021). Rad51 filament dynamics and its antagonistic modulators. Semin Cell Dev Biol 113, 3–13. 10.1016/j.semcdb.2020.06.012.

19. Sun, Y., McCorvie, T.J., Yates, L.A., and Zhang, X. (2020). Structural basis of homologous recombination. Cell Mol Life Sci 77, 3–18. 10.1007/s00018-019-03365-1.

20. Luo, S.C., Yeh, M.C., Lien, Y.H., Yeh, H.Y., Siao, H.L., Tu, I.P., Chi, P., and Ho, M.C. (2023). A RAD51-ADP double filament structure unveils the mechanism of filament dynamics in homologous recombination. Nat Commun 14, 4993. 10.1038/s41467-023-40672-5.

21. Shioi, T., Hatazawa, S., Oya, E., Hosoya, N., Kobayashi, W., Ogasawara, M., Kobayashi, T., Takizawa, Y., and Kurumizaka, H. (2024). Cryo-EM structures of RAD51 assembled on nucleosomes containing a DSB site. Nature 628, 212–220. 10.1038/s41586-024-07196-4.

22. Liu, S., Mine-Hattab, J., Villemeur, M., Guerois, R., Pinholt, H.D., Mirny, L.A., and Taddei, A. (2023). In vivo tracking of functionally tagged Rad51 unveils a robust strategy of homology search. Nat Struct Mol Biol 30, 1582–1591. 10.1038/s41594-023-01065-w.

23. Haas, K.T., Lee, M., Esposito, A., and Venkitaraman, A.R. (2018). Single-molecule localization microscopy reveals molecular transactions during RAD51 filament assembly at cellular DNA damage sites. Nucleic Acids Res 46, 2398–2416. 10.1093/nar/gkx1303.

24. Sharma, N., Coticchio, G., Borini, A., Tachibana, K., Nasmyth, K.A., and Schuh, M. (2024). Changes in DNA repair compartments and cohesin loss promote DNA damage accumulation in aged oocytes. Curr Biol 34, 5131–5148 e5136. 10.1016/j.cub.2024.09.040.

25. Stanyte, R., Nuebler, J., Blaukopf, C., Hoefler, R., Stocsits, R., Peters, J.M., and Gerlich, D.W. (2018). Dynamics of sister chromatid resolution during cell cycle progression. J Cell Biol 217, 1985–2004. 10.1083/jcb.201801157.

26. Ochs, F., Green, C., Szczurek, A.T., Pytowski, L., Kolesnikova, S., Brown, J., Gerlich, D.W., Buckle, V., Schermelleh, L., and Nasmyth, K.A. (2024). Sister chromatid cohesion is mediated by individual cohesin complexes. Science 383, 1122–1130. 10.1126/science.adl4606.

27. Mason, J.M., Dusad, K., Wright, W.D., Grubb, J., Budke, B., Heyer, W.D., Connell, P.P., Weichselbaum, R.R., and Bishop, D.K. (2015). RAD54 family translocases counter genotoxic effects of RAD51 in human tumor cells. Nucleic Acids Res 43, 3180–3196. 10.1093/nar/gkv175.

28. Mazina, O.M., and Mazin, A.V. (2004). Human Rad54 protein stimulates DNA strand exchange activity of hRad51 protein in the presence of Ca2+. J Biol Chem 279, 52042–52051. 10.1074/jbc.M410244200.

29. Bugreev, D.V., Mazina, O.M., and Mazin, A.V. (2006). Rad54 protein promotes branch migration of Holliday junctions. Nature 442, 590–593. 10.1038/nature04889.

30. Spies, J., Waizenegger, A., Barton, O., Surder, M., Wright, W.D., Heyer, W.D., and Lobrich, M. (2016). Nek1 Regulates Rad54 to Orchestrate Homologous Recombination and Replication Fork Stability. Mol Cell 62, 903–917. 10.1016/j.molcel.2016.04.032.

31. Koob, L., Friskes, A., van Bergen, L., Feringa, F.M., van den Broek, B., Koeleman, E.S., van Beek, E., Schubert, M., Blomen, V.A., Brummelkamp, T.R., et al. (2023). MND1 enables homologous recombination in somatic cells primarily outside the context of replication. 1–20. 10.1002/1878-0261.13448.

32. Sakaue-Sawano, A., Kurokawa, H., Morimura, T., Hanyu, A., Hama, H., Osawa, H., Kashiwagi, S., Fukami, K., Miyata, T., Miyoshi, H., et al. (2008). Visualizing spatiotemporal dynamics of multicellular cell-cycle progression. Cell 132, 487–498. 10.1016/j.cell.2007.12.033.

33. Ochs, F., Somyajit, K., Altmeyer, M., Rask, M.B., Lukas, J., and Lukas, C. (2016). 53BP1 fosters fidelity of homology-directed DNA repair. Nat Struct Mol Biol 23, 714–721. 10.1038/nsmb.3251.

34. Spies, J., Lukas, C., Somyajit, K., Rask, M.B., Lukas, J., and Neelsen, K.J. (2019). 53BP1 nuclear bodies enforce replication timing at under-replicated DNA to limit heritable DNA damage. Nat Cell Biol 21, 487–497. 10.1038/s41556-019-0293-6.

35. Strom, L., Lindroos, H.B., Shirahige, K., and Sjogren, C. (2004). Postreplicative recruitment of cohesin to double-strand breaks is required for DNA repair. Mol Cell 16, 1003–1015. 10.1016/j.molcel.2004.11.026.

36. Noda, S., Akanuma, G., Keyamura, K., and Hishida, T. (2023). RecN spatially and temporally controls RecA-mediated repair of DNA double-strand breaks. J Biol Chem 299, 105466. 10.1016/j.jbc.2023.105466.

37. Natsume, T., Kiyomitsu, T., Saga, Y., and Kanemaki, M.T. (2016). Rapid Protein Depletion in Human Cells by Auxin-Inducible Degron Tagging with Short Homology Donors. Cell Rep 15, 210–218. 10.1016/j.celrep.2016.03.001.

38. Lieberman-Aiden, E., van Berkum, N.L., Williams, L., Imakaev, M., Ragoczy, T., Telling, A., Amit, I., Lajoie, B.R., Sabo, P.J., Dorschner, M.O., et al. (2009). Comprehensive mapping of long-range interactions reveals folding principles of the human genome. Science 326, 289–293. 10.1126/science.1181369.

39. Arnould, C., Rocher, V., Finoux, A.L., Clouaire, T., Li, K., Zhou, F., Caron, P., Mangeot, P.E., Ricci, E.P., Mourad, R., et al. (2021). Loop extrusion as a mechanism for formation of DNA damage repair foci. Nature 590, 660–665. 10.1038/s41586-021-03193-z.

40. Yesbolatova, A., Saito, Y., Kitamoto, N., Makino-Itou, H., Ajima, R., Nakano, R., Nakaoka, H., Fukui, K., Gamo, K., Tominari, Y., et al. (2020). The auxin-inducible degron 2 technology provides sharp degradation control in yeast, mammalian cells, and mice. Nat Commun 11, 5701. 10.1038/s41467-020-19532-z.

41. Dumont, A., Mendiboure, N., Savocco, J., Anani, L., Moreau, P., Thierry, A., Modolo, L., Jost, D., and Piazza, A. (2024). Mechanism of homology search expansion during recombinational DNA break repair in Saccharomyces cerevisiae. Mol Cell 84, 3237–3253 e3236. 10.1016/j.molcel.2024.08.003.

42. Ma, C.J., Gibb, B., Kwon, Y., Sung, P., and Greene, E.C. (2017). Protein dynamics of human RPA and RAD51 on ssDNA during assembly and disassembly of the RAD51 filament. Nucleic Acids Res 45, 749–761. 10.1093/nar/gkw1125.

43. Tsabar, M., Eapen, V.V., Mason, J.M., Memisoglu, G., Waterman, D.P., Long, M.J., Bishop, D.K., and Haber, J.E. (2015). Caffeine impairs resection during DNA break repair by reducing the levels of nucleases Sae2 and Dna2. Nucleic Acids Res 43, 6889–6901. 10.1093/nar/gkv520.

44. Ochs, F., Karemore, G., Miron, E., Brown, J., Sedlackova, H., Rask, M.B., Lampe, M., Buckle, V., Schermelleh, L., Lukas, J., and Lukas, C. (2019). Stabilization of chromatin topology safeguards genome integrity. Nature 574, 571–574. 10.1038/s41586-019-1659-4.

45. Eggler, A.L., Inman, R.B., and Cox, M.M. (2002). The Rad51-dependent pairing of long DNA substrates is stabilized by replication protein A. J Biol Chem 277, 39280–39288. 10.1074/jbc.M204328200.

46. Carreira, R., Lama-Diaz, T., Crugeiras, M., Aguado, F.J., Sebesta, M., Krejci, L., and Blanco, M.G. (2024). Concurrent D-loop cleavage by Mus81 and Yen1 yields half-crossover precursors. Nucleic Acids Res 52, 7012–7030. 10.1093/nar/gkae453.

47. Ding, J., Li, X., Shen, J., Zhao, Y., Zhong, S., Lai, L., Niu, H., and Qi, Z. (2023). ssDNA accessibility of Rad51 is regulated by orchestrating multiple RPA dynamics. Nat Commun 14, 3864. 10.1038/s41467-023-39579-y.

48. Chen, H., Lisby, M., and Symington, L.S. (2013). RPA coordinates DNA end resection and prevents formation of DNA hairpins. Mol Cell 50, 589–600. 10.1016/j.molcel.2013.04.032.

49. Piazza, A., Bordelet, H., Dumont, A., Thierry, A., Savocco, J., Girard, F., and Koszul, R. (2021). Cohesin regulates homology search during recombinational DNA repair. Nature Cell Biology 23, 1176–1186. 10.1038/s41556-021-00783-x.

50. Lobachev, K., Vitriol, E., Stemple, J., Resnick, M.A., and Bloom, K. (2004). Chromosome fragmentation after induction of a double-strand break is an active process prevented by the RMX repair complex. Curr Biol 14, 2107–2112. 10.1016/j.cub.2004.11.051.

51. Nakai, W., Westmoreland, J., Yeh, E., Bloom, K., and Resnick, M.A. (2011). Chromosome integrity at a double-strand break requires exonuclease 1 and MRX. DNA Repair (Amst) 10, 102–110. 10.1016/j.dnarep.2010.10.004.

52. Phipps, J., Toulouze, M., Ducrot, C., Costa, R., Brocas, C., and Dubrana, K. (2024). Cohesin complex oligomerization maintains end-tethering at DNA double-strand breaks. Nat Cell Biol. 10.1038/s41556-024-01552-2.

53. Brinkman, E.K., Chen, T., Amendola, M., and Van Steensel, B. (2014). Easy quantitative assessment of genome editing by sequence trace decomposition. Nucleic Acids Research 42. 10.1093/nar/gku936.

54. Haase, R., Royer, L.A., Steinbach, P., Schmidt, D., Dibrov, A., Schmidt, U., Weigert, M., Maghelli, N., Tomancak, P., Jug, F., and Myers, E.W. (2020). CLIJ: GPU-accelerated image processing for everyone. Nature Methods 17, 5–6. 10.1038/s41592-019-0650-1.

55. Uwe Schmidt, M.W., Coleman Broaddus, Gene Myers, AF Frangi, JA Schnabel, C Davatzikos (2018). Medical image computing and computer assisted intervention– MICCAI 2018. 21st International Conference, Granada, Spain, 265–273.

56. Nahidiazar, L., Kreft, M., van den Broek, B., Secades, P., Manders, E.M., Sonnenberg, A., and Jalink, K. (2015). The molecular architecture of hemidesmosomes, as revealed with super-resolution microscopy. J Cell Sci 128, 3714–3719. 10.1242/jcs.171892.

57. Nahidiazar, L., Agronskaia, A.V., Broertjes, J., van den Broek, B., and Jalink, K. (2016). Optimizing Imaging Conditions for Demanding Multi-Color Super Resolution Localization Microscopy. PLoS One 11, e0158884. 10.1371/journal.pone.0158884.

58. Ovesny, M., Krizek, P., Borkovec, J., Svindrych, Z., and Hagen, G.M. (2014). ThunderSTORM: a comprehensive ImageJ plug-in for PALM and STORM data analysis and super-resolution imaging. Bioinformatics 30, 2389–2390. 10.1093/bioinformatics/btu202.

59. Hoogendoorn, E., Crosby, K.C., Leyton-Puig, D., Breedijk, R.M., Jalink, K., Gadella, T.W., and Postma, M. (2014). The fidelity of stochastic single-molecule super-resolution reconstructions critically depends upon robust background estimation. Sci Rep 4, 3854. 10.1038/srep03854.

