## Supplemental figures for "Visualizing homology search in living cells"

Supplemental figure 1: RAD51 forms extended filaments at sites of DNA damage

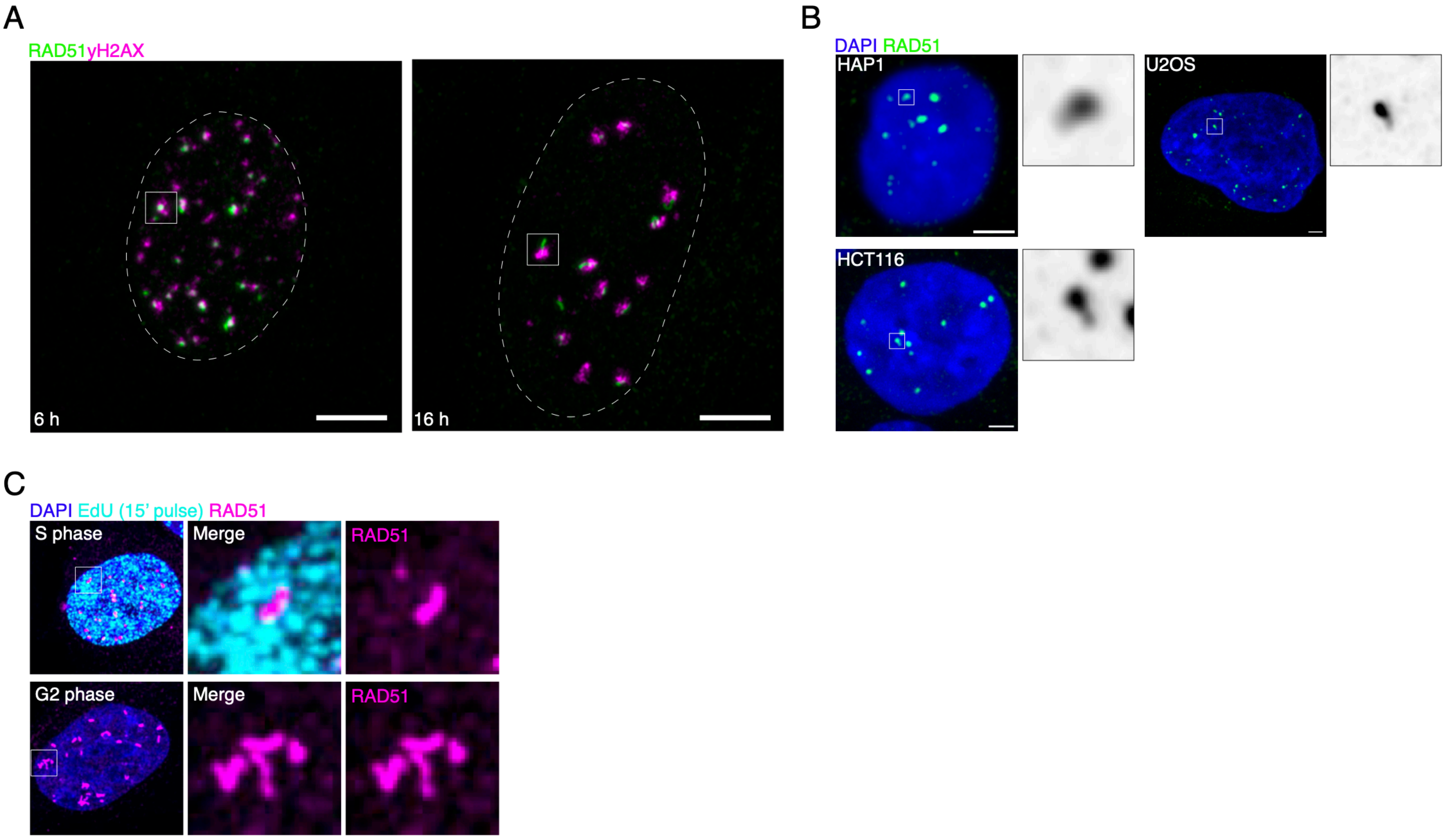

Supplemental figure 2: RAD51 forms extended filaments at sites of DNA damage

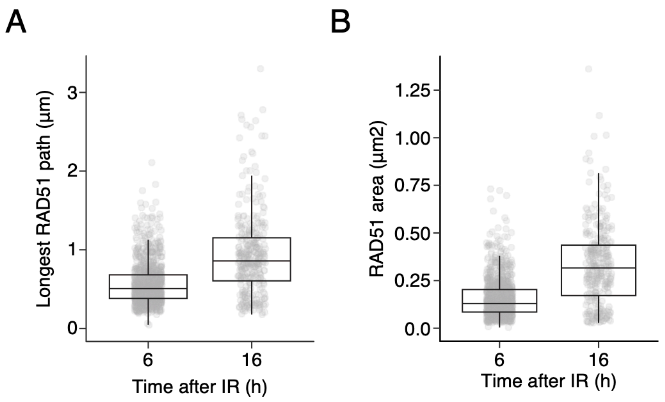

### Supplemental figure 3: RAD54L is required for the resolution of RAD51 filaments

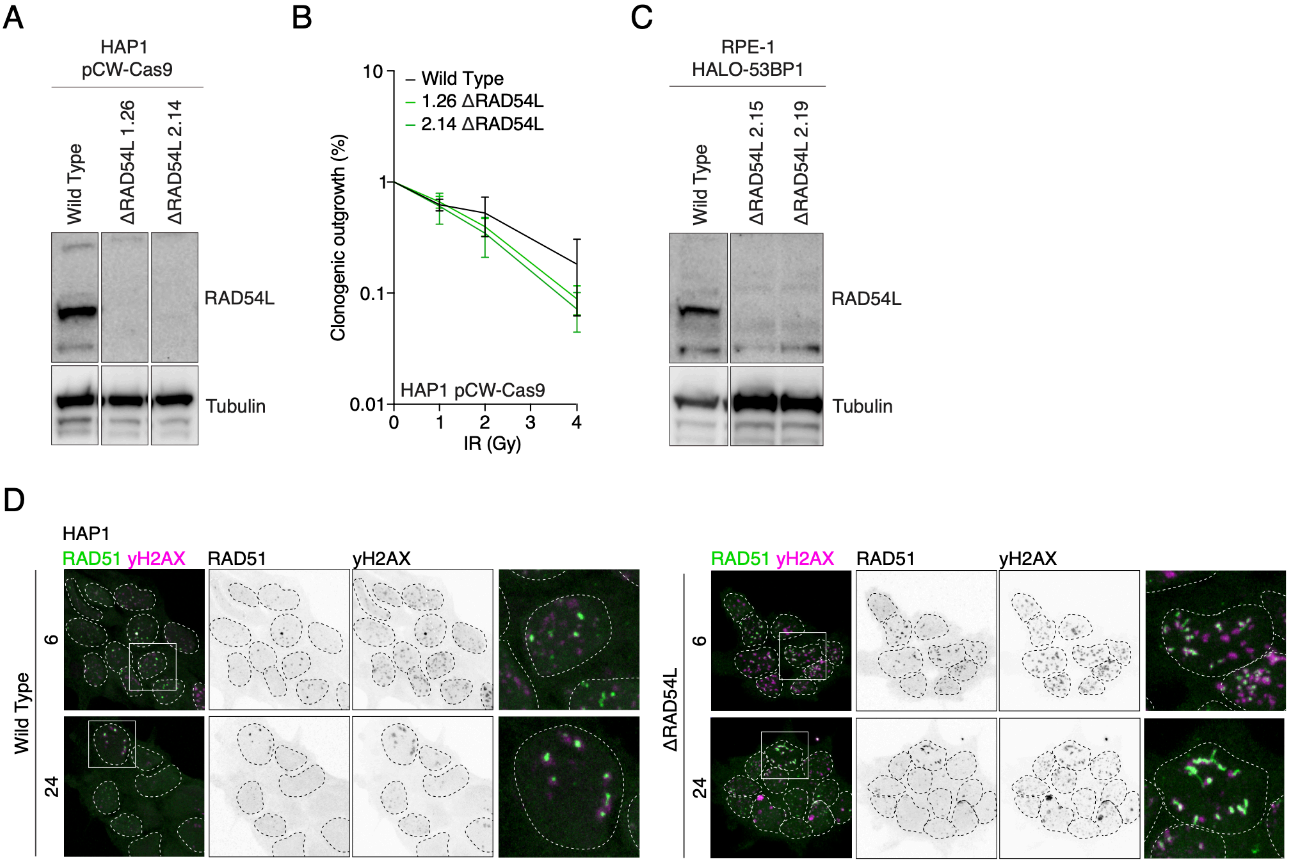

Supplemental figure 4: Two homology search modes: coordinated and independent

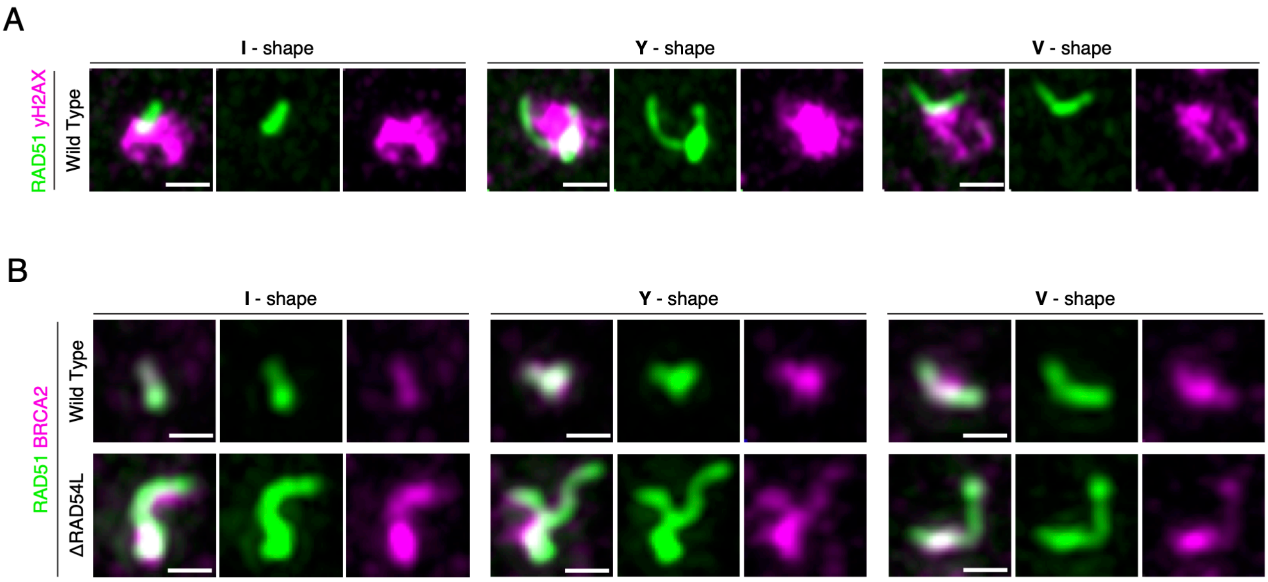

Supplemental figure 5: RAD51 filaments recruit MND1

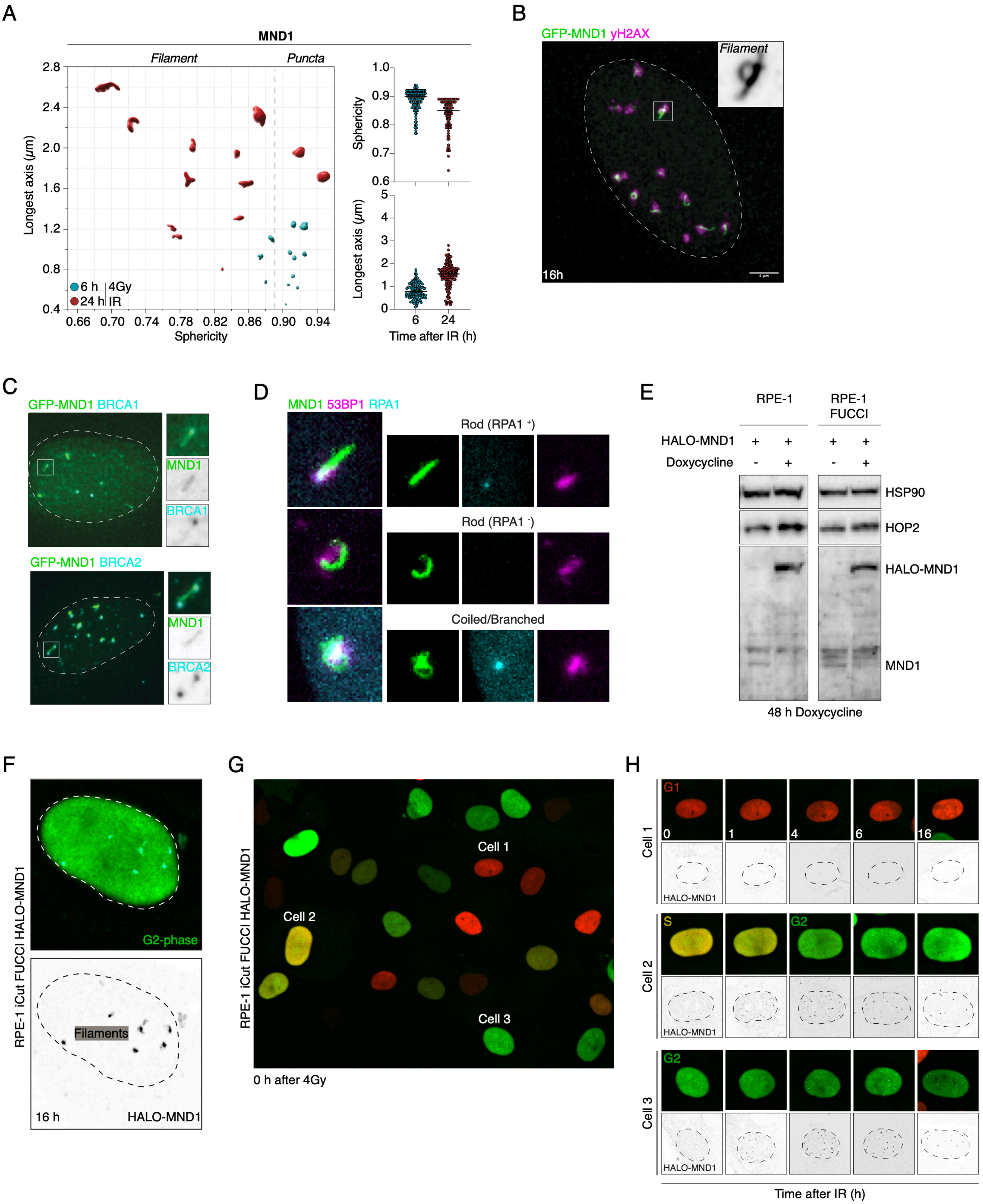

Supplemental figure 6: The capture event in homology search

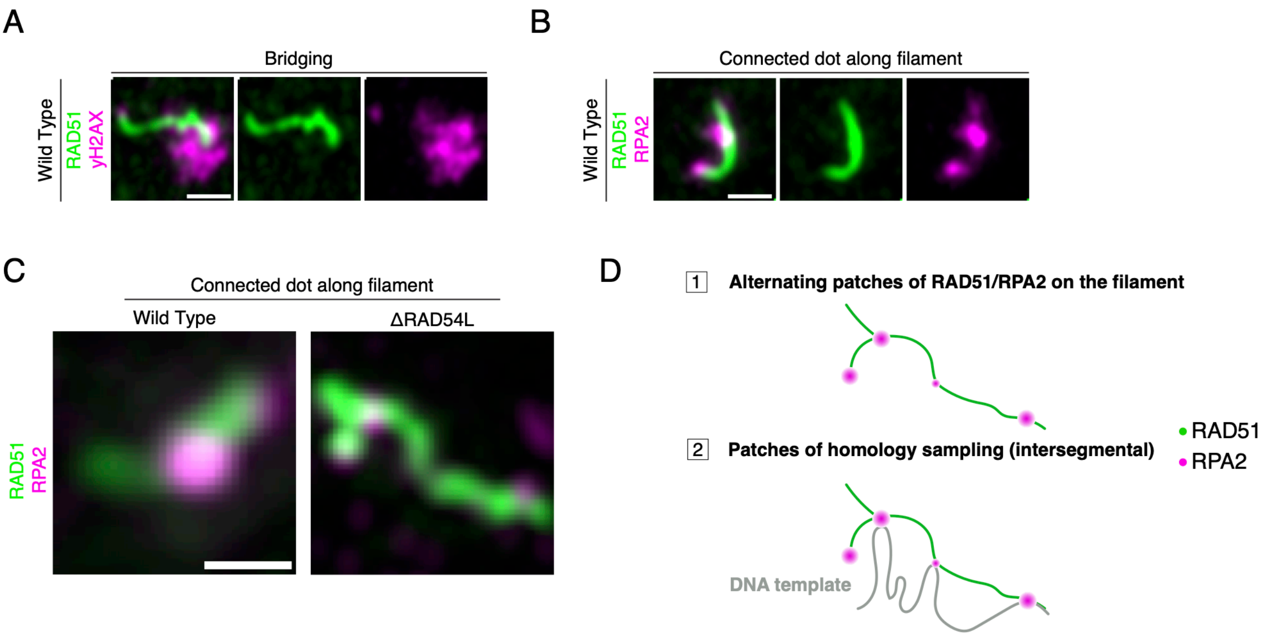

Supplemental figure 7: Cohesin, not CTCF, controls RAD51-mediated repair

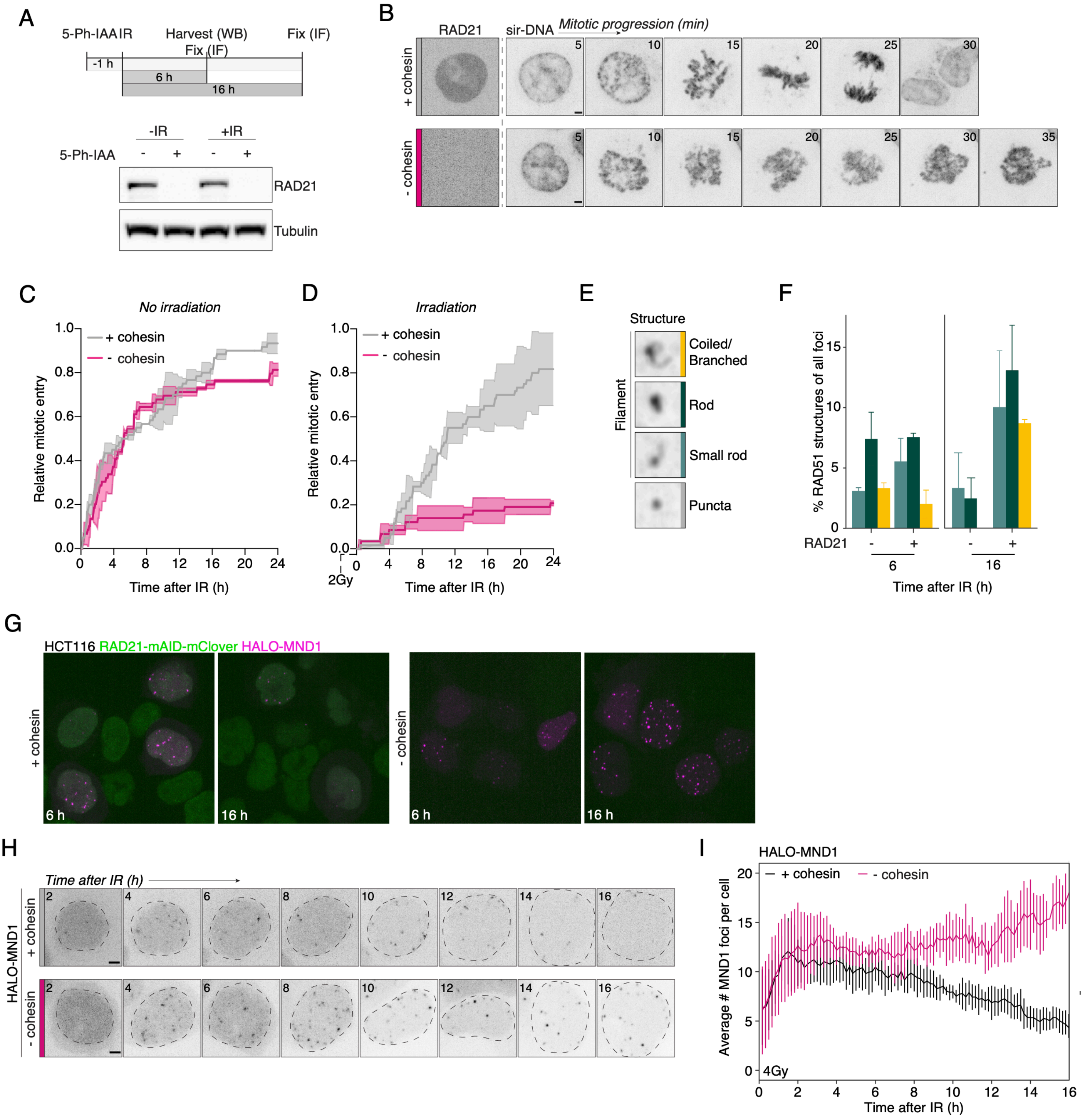

### Supplemental figure 8: Cohesin, not CTCF, controls RAD51-mediated repair

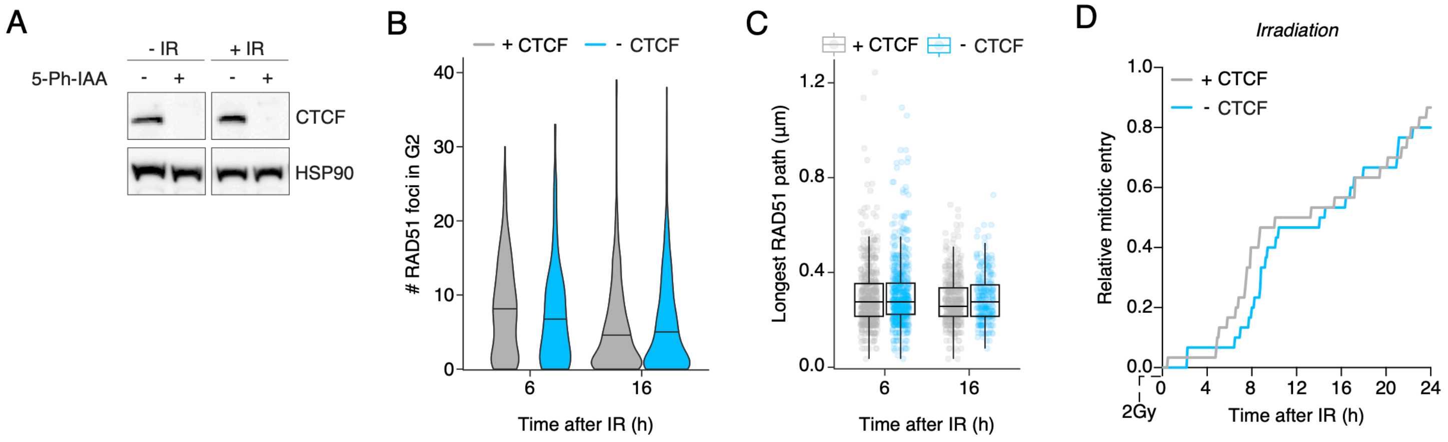

Supplemental figure 9: Double act: RAD54L and cohesin unite for effective homology search

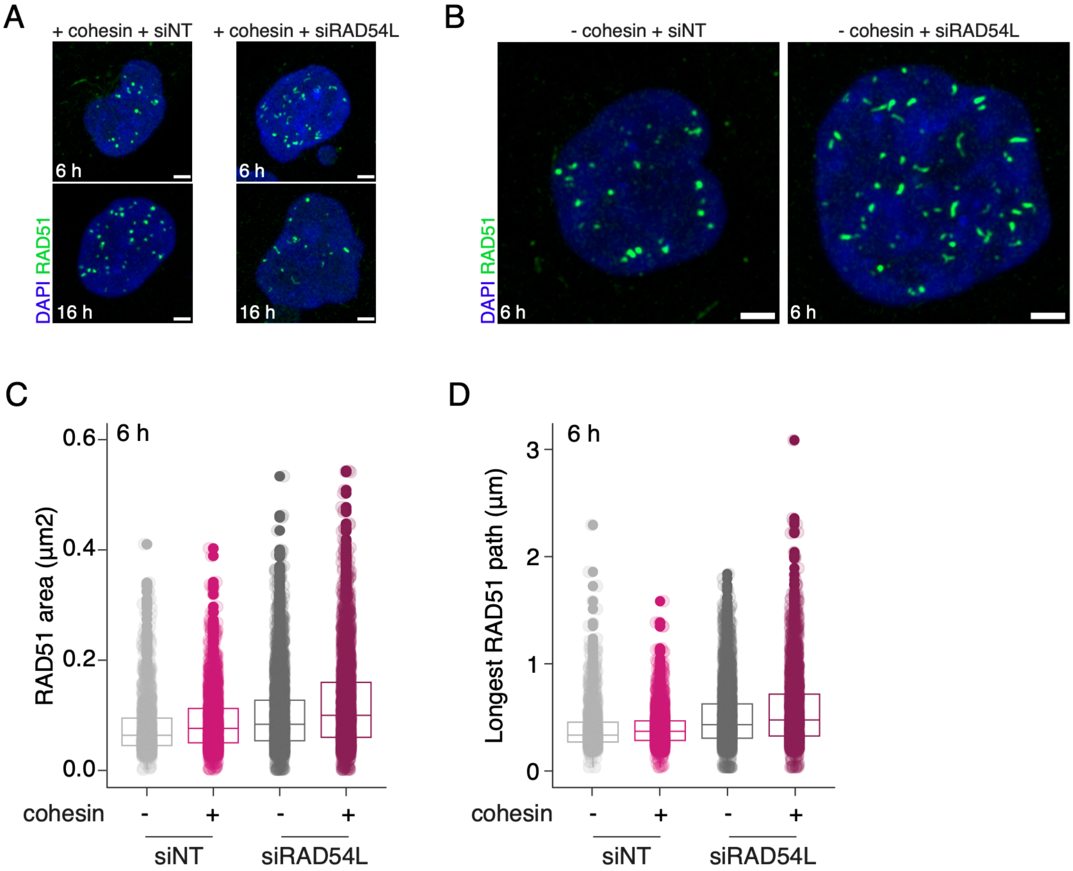
